# Single-cell RNA Sequencing Reveals the Temporal Diversity and Dynamics of Cardiac Immunity after Myocardial Infarction

**DOI:** 10.1101/2021.07.19.452902

**Authors:** Kaiyu Jin, Shan Gao, Penghui Yang, Rongfang Guo, Dan Li, Yunsha Zhang, Xiaoyan Lu, Guanwei Fan, Xiaohui Fan

**Affiliations:** Pharmaceutical Informatics Institute, College of Pharmaceutical Sciences, Zhejiang University, Hangzhou 310058, China; First Teaching Hospital of Tianjin University of Traditional Chinese Medicine, Tianjin Key Laboratory of Translational Research of TCM Prescription and Syndrome, Tianjin 300193, China; State Key Laboratory of Component-Based Chinese Medicine, Tianjin 301617, China; School of Integrative Medicine, Tianjin University of Traditional Chinese Medicine, Tianjin 300193, China; iMedicine Lab, Alibaba-Zhejiang University Joint Research Center of Future Digital Healthcare, Hangzhou 310058, China

**Keywords:** myocardial infarction, temporal dynamics, cardiac immune populations, macrophages, single-cell RNA sequencing

## Abstract

Myocardial infarction (MI) is strongly associated with the temporal regulation of cardiac immunity. However, a variety of current clinical trials have failed because of the lack of post-MI immunomodulating/anti-inflammatory targets. We performed single-cell RNA sequencing analysis of cardiac *Cd45*^+^ immune cell at 0, 3, 7, and 14 days after injury in a mouse left anterior descending coronary artery ligation model. Major immune cell populations, distinct subsets, and dynamic changes were identified. Macrophages (Mø)/monocytes were most abundant, peaking at 3 days after infarction. Mø-5 and Mø-6 were the predominant infiltrated subsets at this time point, with strong expression of inflammatory factors. Further analysis demonstrated that suppressing these sets attenuated pathological MI progression by preventing subsequent leukocyte extravasation and adverse remodeling. We also detected abundant apoptotic neutrophils and a profibrotic macrophage subset on days 7 and 14 respectively. These results provide a basis for developing cell type- and time-specific interventions in MI.

**Graphic abstract:** 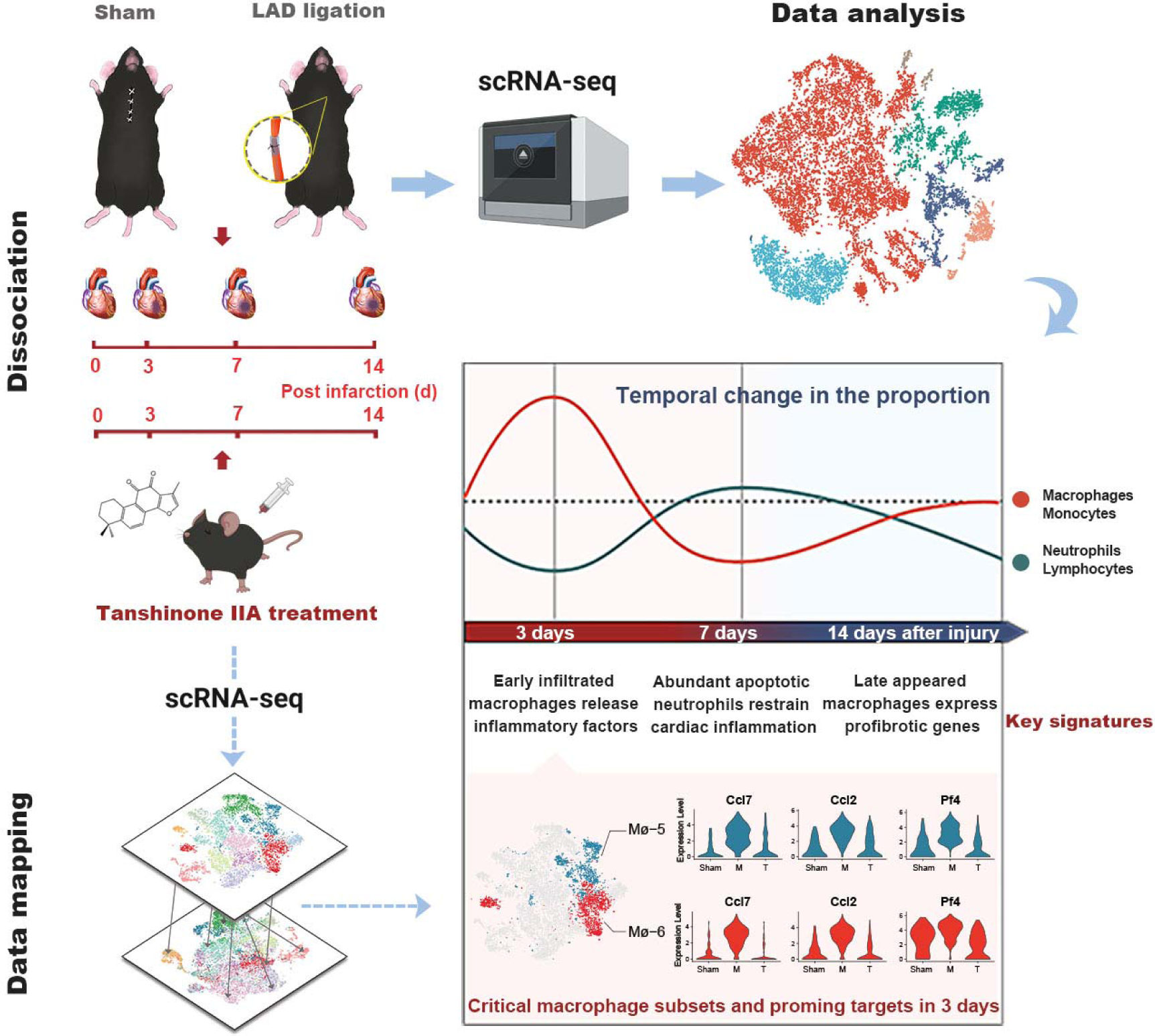

Temporal regulation of cardiac immunity is strongly associated with the onset, progression, and outcomes of MI. Our study of time-series scRNA-seq on the whole cardiac immune cells isolated from the LAD ligation mice and the tanshinone IIA treated mice shed light on the underlying pathology of MI.

## 1. Introduction

Myocardial infarction (MI), a significant cause of mortality and morbidity worldwide,^[1]^ is characterized by the recruitment and activation of immune cells. These cells orchestrate very complex immune responses that can either propagate or defend against heart disease. For example, immune cells are involved in post-MI initial pro-inflammatory processes as well as later anti-inflammatory responses, including by playing roles in wound healing or remodeling.^[2]^ Therefore, cardiac immunity is considered as an essential pathophysiologic factor and target in the context of MI. However, numerous clinical trials of post-MI immunomodulating/anti-inflammatory targets have shown inconclusive and controversial results.^[3]^ Hence, studies are needed to understand what, how, and in what time frame the immune system is activated, providing a basis for modulating the immune-inflammatory mechanisms underlying the myocardial healing process.

Cardiac immune cells are highly heterogeneous, and their activation in the progression of MI is distinct.^[4]^ Some cell types such as macrophages and monocytes, which play a central role in multiple pathophysiological processes of MI, are comprised of diverse subsets because of their different origins and functions.^[5]^ However, much less is known about the complex diversity of cardiac immunity. Recently, an innovative high-throughput technology, single-cell RNA sequencing (scRNA-seq), has increased our understanding of the major immune cell types in the mammalian heart and their dynamics during disease progression. Dick *et al*^[6]^ identified functionally distinct cardiac macrophage subsets that are hierarchically replaced by monocytes after infarction. Vafadarnejad *et al*^[7]^ revealed temporal diversity and local transition of neutrophils within the ischemic tissue. Moreover, Farbehi *et al*^[8]^ described the heterogeneity and dynamics of several cardiac immune cell populations simultaneously by performing scRNA-seq of the total non-cardiomyocyte fraction. However, this study detected a relatively lower quantity of immune cells and did not evaluate later stages, which may be more strongly associated with advanced cardiac remodeling. Therefore, systematic analyses of temporal dynamics of cardiac immunity following MI are needed to develop time-specific immunomodulatory interventions.

Here, we investigated the comprehensive cardiac immune landscape by analyzing the high-throughput single-cell transcriptomic profiles of 16,380 mouse cardiac *Cd45*^+^ cells in a longer time window after MI. We identified major immune cell populations as well as their distinct subsets based on unique molecular signatures throughout the disease. We focused predominately on macrophages, which expanded dramatically early post-MI and displayed critical roles in either promoting an excessive inflammatory response or providing tissue repair. We evaluated the gene profiles during disease progression and characterized the stepwise changes and transitions in cardiac macrophages. We also determine the timing and roles of neutrophil and lymphocyte populations. Additionally, detrimental macrophage subsets showing potential as therapeutic targets were identified by performing parallel scRNA-seq on tanshinone IIA treated murine hearts. The detailed molecular signatures identified in our study improve the understanding of the functions, signaling pathways, and interactions within distinct immune subtypes and will facilitate identification of specific features that can be used as prognostic and therapeutic targets in human MI.

## 2. Results

### 2.1 Temporal immune composition in the heart by single-cell RNA sequencing after MI

To identify diverse cardiac immune cell types and comprehensively characterize their dynamic alterations during the pathological progression of MI, we performed scRNA-seq analysis of sorted *Cd45*^+^ cells from the murine heart at different time points (0, 3, 7, and 14 days) after MI using the 10X Genomics platform (**Figure 1A**). These four time points reflected the major stages in MI development, corresponding to a progressive increase in inflammatory cell infiltration and myocardial fibrosis (**Figure 1B**; **Figure S1 and S2A**). Thus, we covered the early acute inflammatory stage, intermediate repairing stage, and advanced remodeling stage. We pooled 17,932 individual cells from all four-time points into a single dataset after quality trimming and filtering (**Figure S3A and Table S1**). The origin of each cell was determined (as visualized by different colors in **Figure S3C**). Additionally, to evaluate the biological differences between stages, we calculated the correlations in gene expression levels between samples, implying that all time points were quite similar (**Figure S4A**).

**Figure 1.**
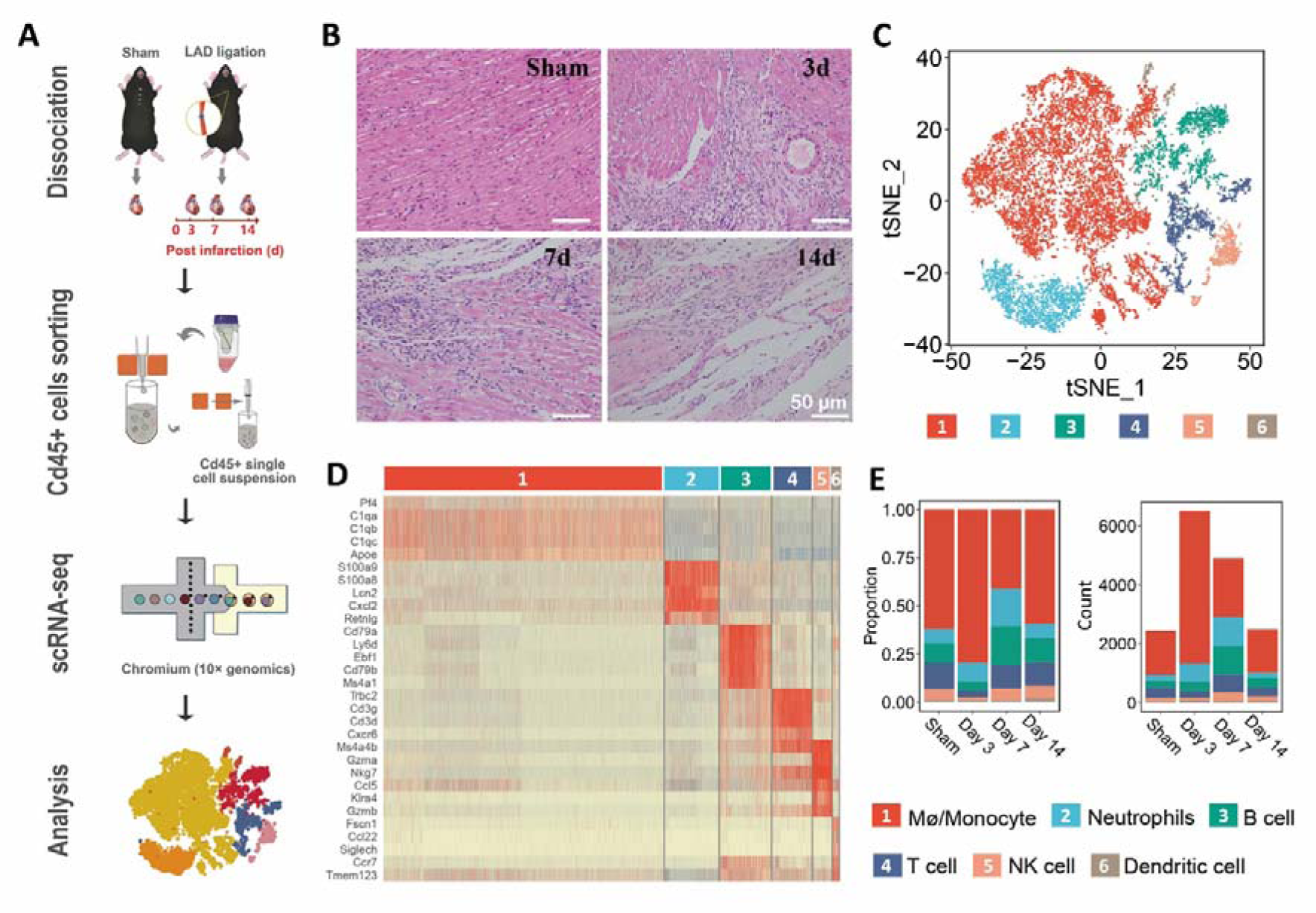
Single-cell RNA sequencing of cardiac *Cd45*^+^ cells from four samples in the MI dataset reveals the post-infarction dynamics of six immune cell populations. A. Schematic overview of the 10X chromium scRNA-seq procedure used to evaluate *Cd45*^+^ immune cells isolated from the hearts of C57/BL mice 0, 3, 7, and 14 days after left anterior descending artery ligation. Five mice at each time point were processed in parallel. B. Representative HE staining images of hearts from the sham group and different MI groups. Compared with the sham group, severe injuries were observed in the MI groups, including abnormal myocardial morphology, inflammatory cell infiltration, and aggravated fibrosis. C. tSNE plot of 16,380 cells pooled from the sham and three MI groups, showing the immune cell populations identified based on known cell type markers. D. Heatmap of top 5 differentially expressed genes in each subpopulation. E. Fraction of immune cell populations relative to all cardiac immune cells as a function of disease progression (left). Cell numbers of each population were determined by unique molecular identifiers (UMIs) (right)

Unsupervised clustering and t-distributed stochastic neighbor embedding (t-SNE) dimensionality reduction implemented in the Seurat package was performed on this complex dataset (**Figure 1C**). Six major immune cell populations, including macrophage (Mø)/monocyte, neutrophil, T cell, B cell, natural killer cell (NK cell), and dendritic cell populations, were identified based on the expression of highly variable genes (**Figure 1D; Figure S4B and Table S2**). Cardiac immune cell heterogeneity already existed under homeostatic conditions; however, the Mø/monocyte population was most abundant, accounting for more than half of the total cells (**Figure 1E**). Moreover, this population exhibited dramatic expansion on day 3 after MI (**Figure 1E**), as validated by flow cytometry (**Figure S2B**). These results highlight the indispensable role of the Mø/monocyte population in orchestrating the initial immune response to injury. Notably, apparent alterations in cluster proportions were observed on day 7 after MI, when T cells, B cells, and neutrophils were all increased substantially (**Figure 1E**). This agrees with the results of previous studies showing that lymphocytes and neutrophils exhibited slower but essential responses.^[2, 9]^ Finally, a comparable immune composition to that in the sham group was observed on day 14 (**Figure 1E**), indicating successful self-control of the disease.

These results suggest that within the studied time window (days 0–14), different immune subpopulations infiltrated the heart at different time points and reacted collectively to MI via distinct functions.

### 2.2 Defining cardiac macrophage subsets with distinct functions at all time points

Macrophages were the most abundant immune cells residing in the heart or invading the infarcted myocardium. Extensive studies have shown that they are a highly heterogeneous population with varying functions; therefore, we performed second-level clustering analyses of cells in this population to better understand their characteristics and potential roles in MI progression. In total, 12 clusters (a minor cluster was excluded because of non-specific gene expression) were identified, each with a unique transcriptional profile and functions (**Figure 2A–B; Excel File S1 and S2**). We first identified monocytes based on well-known conventional markers, such as *Ly6c2*, *Plac8*, and *Hp*, which were absent/sparse in other cells. In contrast, these cells expressed high levels of some prototypical macrophage genes (*C1qa* and *C1qb*)^[10]^ (**Figure S5A**). Next, we examined the expression of the signature genes *Ccr2* and *MHC* C (*H2-Eb1*, *H2-DMb1*, and *H2-DMa*), which indicate a monocytic origin and robust inflammatory potential.^[11]^ Mø-1, Mø-2, Mø-3, and Mø-4 were *MHC*^high^ subsets (**Figure 2C**) and enriched for Gene Ontology (GO) terms related to antigen processing and presentation (**Excel File S2**). Mø-4 was the only subset lacking expression of *Ccr2* (**Figure 2C**) but still shared many inflammatory features (*Cxcl9*, *Gpb2*, *Ly6a*, *Ly6i*, *Cfb*, etc.) with Mø-1 (**Figure 2B**). We observed that Mø-2 and Mø-3 had expanded by day 3 after MI, whereas Mø-4 was more noticeable at a later stage (**Figure 2D**). In summary, these *MHC*^high^ subsets mainly displayed pro-inflammatory functions and varied at different time points of MI progression.

**Figure 2.**
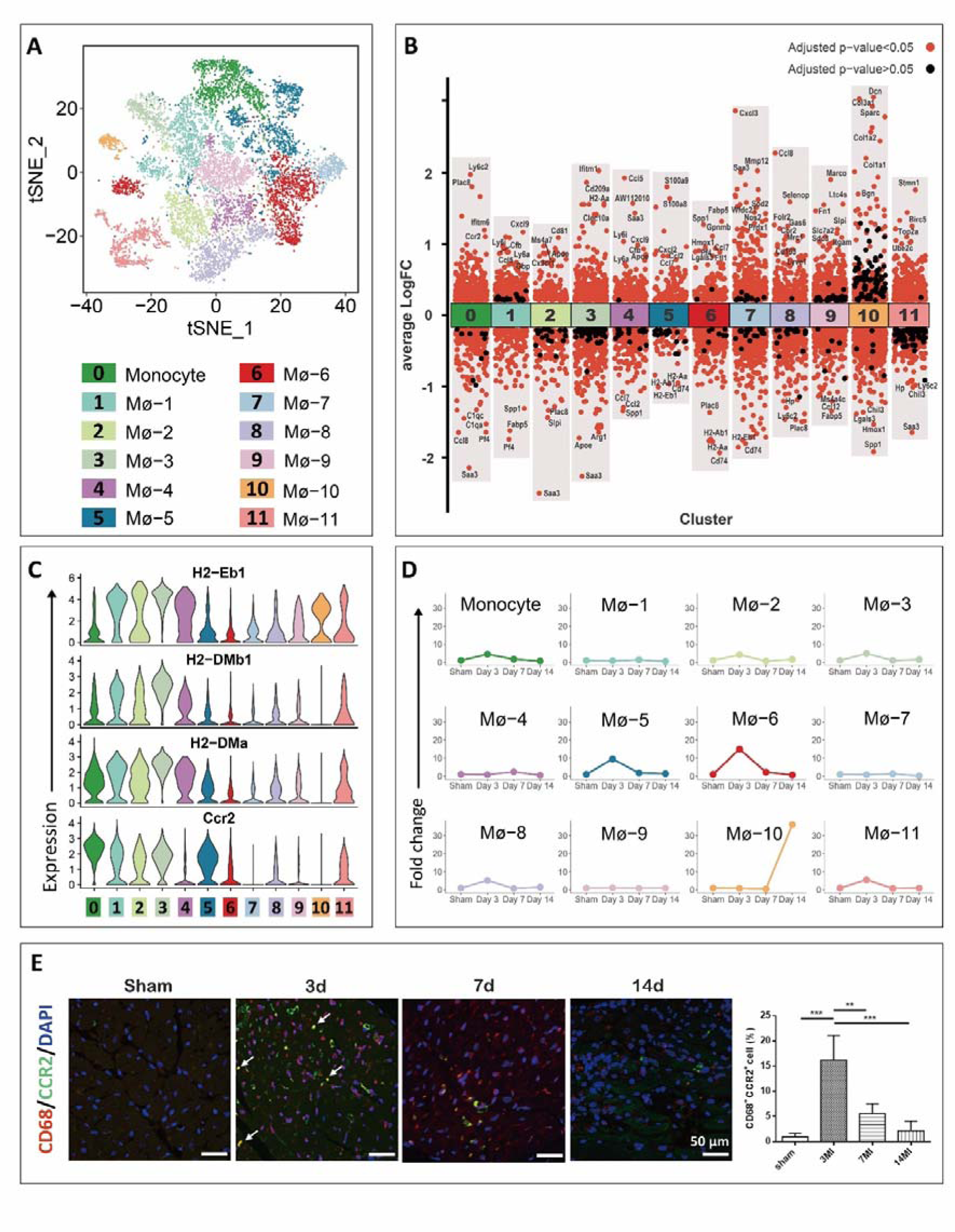
Cardiac macrophages are highly heterogeneous and exhibit distinct functions in MI progression. A. t-SNE plot of 10,216 monocytes and macrophages shows 12 subsets merged from four samples in the MI dataset. All cells were obtained from the Mø/monocyte subpopulation in Fig. 1C and reclustered to increase resolution. B. Differential gene expression analysis showing up- and down-regulated genes across all 12 subsets. The adjusted *P* value < 0.05 is indicated in red, whereas the adjusted *P* value > 0.05 is indicated in black. C. Expression of *MHC* genes (*H2-Eb1*, *H2-DMb*, and *H2-DMa*) and *Ccr2* in each subset. Normalized expression value is indicated on the y-axis D. Relative cell number fold-change (relative to sham) for each subset as a function of progression. E. Representative images of immunostaining for the macrophage marker CD68 (red) and signature genes CCR2 (green) in the murine hearts at 0, 3, 7, and 14 days after MI. Nuclear was counterstained with DAPI (blue). White arrows indicate CCR2^+^CD68^+^ cells in the heart. Quantification of the percentage of CCR2^+^CD68^+^ cells was determined from images using ImageJ software (NIH, Bethesda, MD, USA).

Mø-5 and Mø-6 were both enriched for characteristics related to leukocyte migration and chemotaxis (**Excel File S2**). They showed the up-regulation of genes (*S100a9*, *S100a8*, *Ccl7*, etc.) strongly associated with subsequent leukocyte extravasation (**Figure 2B**). However, these two sets showed different *Ccr2* expression levels and some distinct functions (**Figure 2C; Excel File S2**). Mø-5 expressed a high level of *Ccr2* which was increased dramatically on day 3 (**Figure 2D**), indicating that it was a typical postinfarction inflammatory cluster. In contrast, Mø-6 exhibited lower *Ccr2* expression and displayed more complex functions, including the selective expression of genes with anti-inflammatory potential (*Gpnmb* and *Hmox1*)^[10, 12]^ and genes responsible for iron storage (*Ftl*)^[13]^ (**Figure 2B**). Additionally, these cells showed differential expression of fatty acid (FA) binding protein 5 (*Fabp5*) (**Figure 2B**), which is vital for FA transport from the circulation to the heart, providing cardiac protection via an energy metabolism pathway.^[14]^ Similarly, this set was highly amplified, showing approximately 15-fold expansion from days 0 to 3 (**Figure 2D**). Therefore, this set was also a postinfarction cluster with superimposed characteristics.

The next four subsets showed low levels of *Ccr2* and *MHC* expression (**Figure 2C; Figure S5B**) and were largely tissue-resident macrophages. Mø-7 showed the ability to protect against host cell damage (**Excel File S2**) and expanded to a minor extent by day 7 (**Figure 2D**). In addition, Mø-7 uniquely expressed genes (*Sod2* and *Prdx1*) encoding antioxidant enzymes^[10, 15]^ (**Figure 2B**). Mø-8 expressed classic resident-macrophage markers (*Lyve1*, *Gas6*, and *Cbr2*) and scavenging genes (*Selenop*, *Mrc1*, and *Cd163*)^[16]^ (**Figure 2B**), which were predominately enriched for GO terms related to receptor-mediated endocytosis (**Excel File S2**). As shown in **Figure 2D**, this set greatly increased on day 3. Mø-9 and Mø-10 performed many functions related to tissue integrity and repair; however, Mø-9 expressed numerous genes contributing to angiogenesis, e.g., *Fn1*, *Slc7a2*, and *Sdc3*,^[17]^ whereas the other subset expressed collagen-related genes, e.g., *Col3a1*, *Col1a2*, *Col1a1*, *Dcn*, and *Bgn*,^[18]^ which contribute to tissue fibrosis (**Figure 2B; Excel File S2**). This profibrotic cluster manifestly appeared at 14 days after MI and may be involved in improving cardiac regeneration.^[19]^ Finally, Mø-11 was characterized by high and restricted expression of cell cycle and proliferation markers (e.g., *Stmn1*, *Birc5*, *Top2a*, and *Ube2c*)^[10]^ (**Figure 2B**).

Interestingly, we observed that nearly all *Ccr2*^low^ subsets (Mø-6, Mø-7, Mø-8, and Mø-9) exclusively expressed *Ccr2* on day 3 (**Figure 2C; Figure S5C**), suggesting that inflammatory cells still had precedence at this early phase. Immunofluorescence staining showed a similar result, with a large number of *Ccr2*^high^ macrophages emerging at 3 days after MI (**Figure 2E**).

### 2.3 Dynamic variation in macrophage subsets after MI

We performed pseudotime analyses for trajectory deductions over time to improve the understanding of temporal dynamics and transitions among distinct macrophage subsets. We applied Monocle 3 to construct a developmental trajectory and superimposed Seurat-defined clusters on this trajectory (**Figure 3A; Figure S6A**). The main path along the pseudotime started with monocytes, followed by *MHC*^high^ sets, and finally resident-like or postinfarction sets. Gradient expression of the corresponding marker genes is shown in **Figure S6C**.

**Figure 3.**
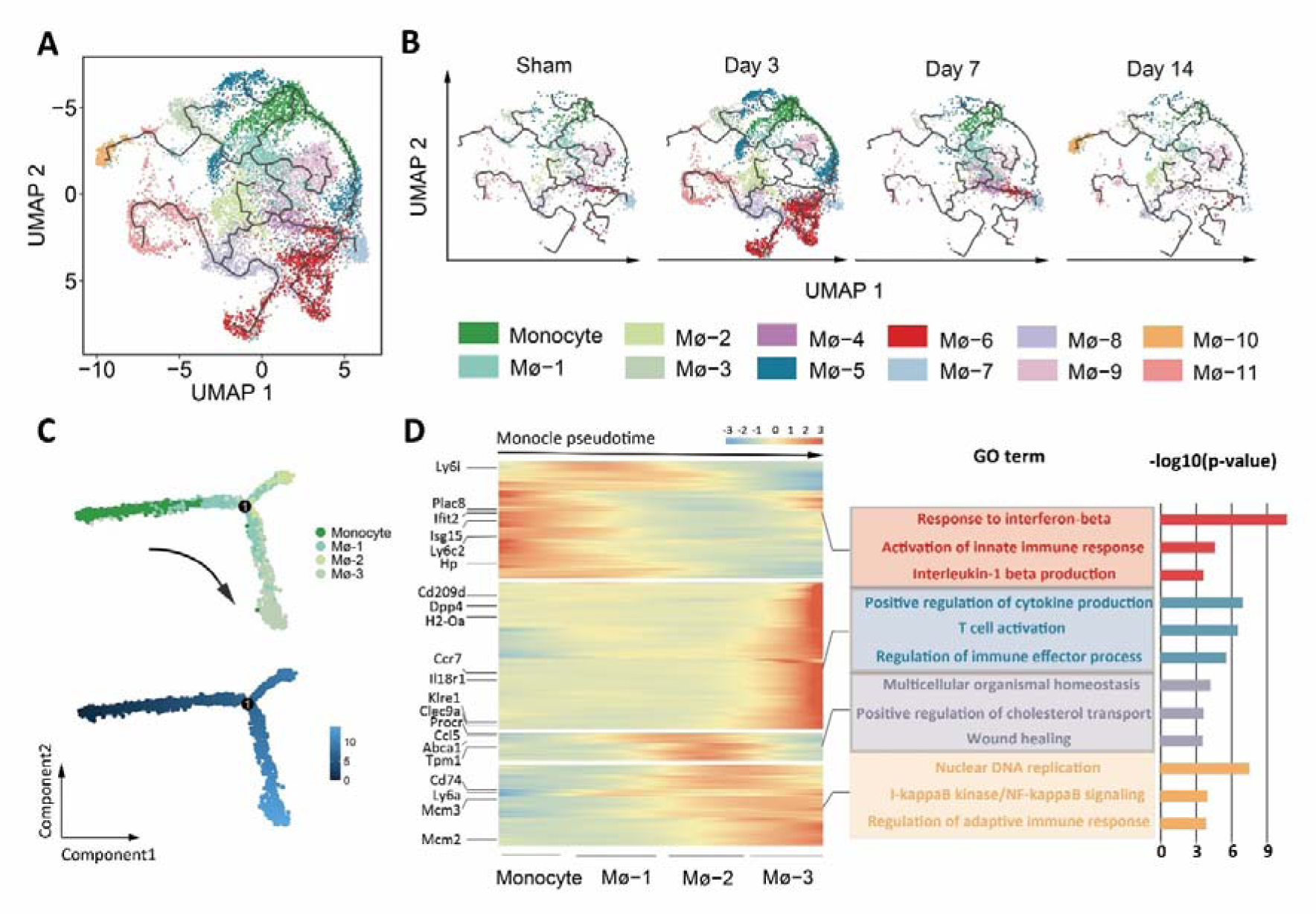
Temporal dynamics of monocytes and macrophages in both real MI progression and pseudotime. A. UMAP visualization of cells arranged along trajectories, colored by identified subset or inferred pseudotime (**Figure S6A**). B. Visualization of cardiac macrophage trajectories split by samples. C. Focused Monocle trajectory analysis including only the monocyte subset, Mø-1 subset, Mø-2 subset, and Mø-3 subset. D. Heatmap of differentially expressed genes, ordered based on their common kinetics through pseudotime using Monocle. The relative position of individual Seurat-based subsets across pseudotime is depicted below. Distinct GO terms and their *P*-value associated with each model are shown on the right.

Mø-5 and Mø-6 were the predominant postinfarction clusters, expanding dramatically at an early stage (3 days) (**Figure 2D**); therefore, we further explored their origins of differentiation. Considering that Mø-5 highly expressed *Ccr2* and appeared along with monocytes in the trajectory (**Figure 3B; Figure S6D**), it may be a typical postinfarction inflammatory cluster directly differentiated from monocytes. Similarly, some macrophages in Mø-6 showed high expression of the monocytic gene *Ccr2* on day 3 (**Figure S6D**). However, this set appeared at the most distant part of the trajectory, and some fraction shared the same branch as Mø-8 (**Figure 3B**), which was the resident-like subset and similarly expanded by day 3. These results indicate that Mø-6 arises from circulating blood cells as well as the self-expanded resident subset.

Additionally, we analyzed the developmental relationships between monocyte and the *MHC*^high^*Ccr2*^high^ subsets at the same stage to clearly distinguish their functions. Mø-1 exhibited a superimposed inflammatory signature and appeared under homeostatic conditions (**Figure 3B**). However, it was among the few subsets showing a reduction after 3 days, along with expansion of the other two sets (**Figure 2D**); therefore, we hypothesized that macrophages in Mø-2 and Mø-3 arise from long-lived Mø-1 rather than from monocytes. This transition was confirmed by restricted Monocle 2 analysis (**Figure 3C**). Predicted gene expression was plotted in pseudotime to track changes across different macrophage states, revealing modules with distinct GO functions (**Figure 3D**). Mø-2 was distinguished by its enrichment in wound healing, consistent with previous observations showing that it expressed more resident macrophage markers (*Apoe*, *Ms4a*, *Cx3cr1*, *Cd81*, etc.)^[10, 16a]^ and fewer inflammatory markers (**Figure 2B**). Mø-3 were enriched in GO terms related to regulation of T cell activation and differed by high expression of *Cd209d*, *Dpp4*, and *H2-O5*.

Collectively, complicated transitions within the Mø/monocyte population were observed by day 3. A very large number of macrophages in Mø-5 and Mø-6 were mainly derived from monocytes. They play an essential role in the profound leukocyte extravasation, which may be detrimental to disease development. Additionally, long-lived macrophages could further differentiate or self-proliferate. These variations were crucial for regulating the injuries. By 7 days, the postinfarction subsets had almost disappeared, whereas Mø-1 and Mø-4 were increased (**Figure 3B; Figure S6B**). They both showed high expression levels of *MHC* genes, corresponding to the coincident adaptive immune response. Inflammatory signatures were still observed at this time point. However, the proportions of the entire subsets were apparently changed (**Figure S6B**). Some clusters with inflammatory phenotype decreased in quantity, suggesting a turning point from the prolongation of inflammation to restraint. Macrophages involved in tissue repair and integrity were expanded, including a collagen-related subset (**Figure S6B**), which are associated with cardiac remodeling.

### 2.4 Identifying phase-specific molecular signatures of macrophages after MI

To further clarify temporal molecular changes within cardiac macrophages in the context of MI, we analyzed differential gene expression at various time points. We performed GO enrichment analysis of the differentially expressed genes to determine the biological features of each stage. In total, 382 genes were up-regulated at 3 days after MI and were enriched in chemokine or cytokine activity (**Figure 4A–B; Figure S7A**). For example, we detected high levels of platelet factor 4 (*Pf4*), chemokine (C-C motif) ligand 2 (*Ccl2*)/MCP-1, and chemokine (C-C motif) ligand 7 (*Ccl7*)/MCP-3 expression on day 3. These chemokines act as potent neutrophil and monocyte attractants, leading to the deterioration of myocardial function and left ventricular remodeling.^[20]^ However, we also found that *Spp1* and *Lgals3* were up-regulated, which tend to induce reparative macrophage polarization and promote fibrosis.^[10, 21]^ In summary, genes expressed on day 3 showed the highest chemokine score and played a critical role in early leukocyte infiltration (**Figure 4C**). The predominant feature of macrophages on day 7 was antigen presentation and processing (**Figure 4B–C; Figure S7B**), which contributed to the adaptive immune response. IFN-γ-induced genes were also upregulated (**Figure 4A**). *Isg15*, which is involved in the IKK/NF-κB pathway, causes an inflammatory response and myocyte atrophy.^[22]^ Its overexpression has a synergetic effect with STAT1 on cardiomyocyte death from acute myocardial ischemia/reperfusion injury.^[23]^ However, we observed up-regulation of antioxidant genes (*Prdx5*, *Sod2*, and *Acod1*) (**Figure 4A**), which are essential for eliminating excessive reactive oxygen species produced after MI (**Figure 4B; Figure S7B**). Additionally, the reactive oxygen species metabolism-related subset Mø-7 expanded by day 7 (**Figure 2D**), indicating that modulation of oxidative stress responses was initiated. At 14 days, a range of genes corresponding to extracellular matrix components was upregulated (**Figure 4A–B; Figure S7C**). Previous studies demonstrated that macrophages express almost all known collagen and collagen-related mRNAs, which not only contribute to the modulation of cell–cell and cell–matrix interactions but also profoundly influence pathophysiological conditions *in vivo*.^[18]^ These results revealed stepwise molecular changes in cardiac macrophages (**Figure 4D**), which were validated by immunofluorescence staining (**Figure 4E**). Characteristic genes representing distinct biological functions at each stage showed time-specific expression during MI progression.

**Figure 4.**
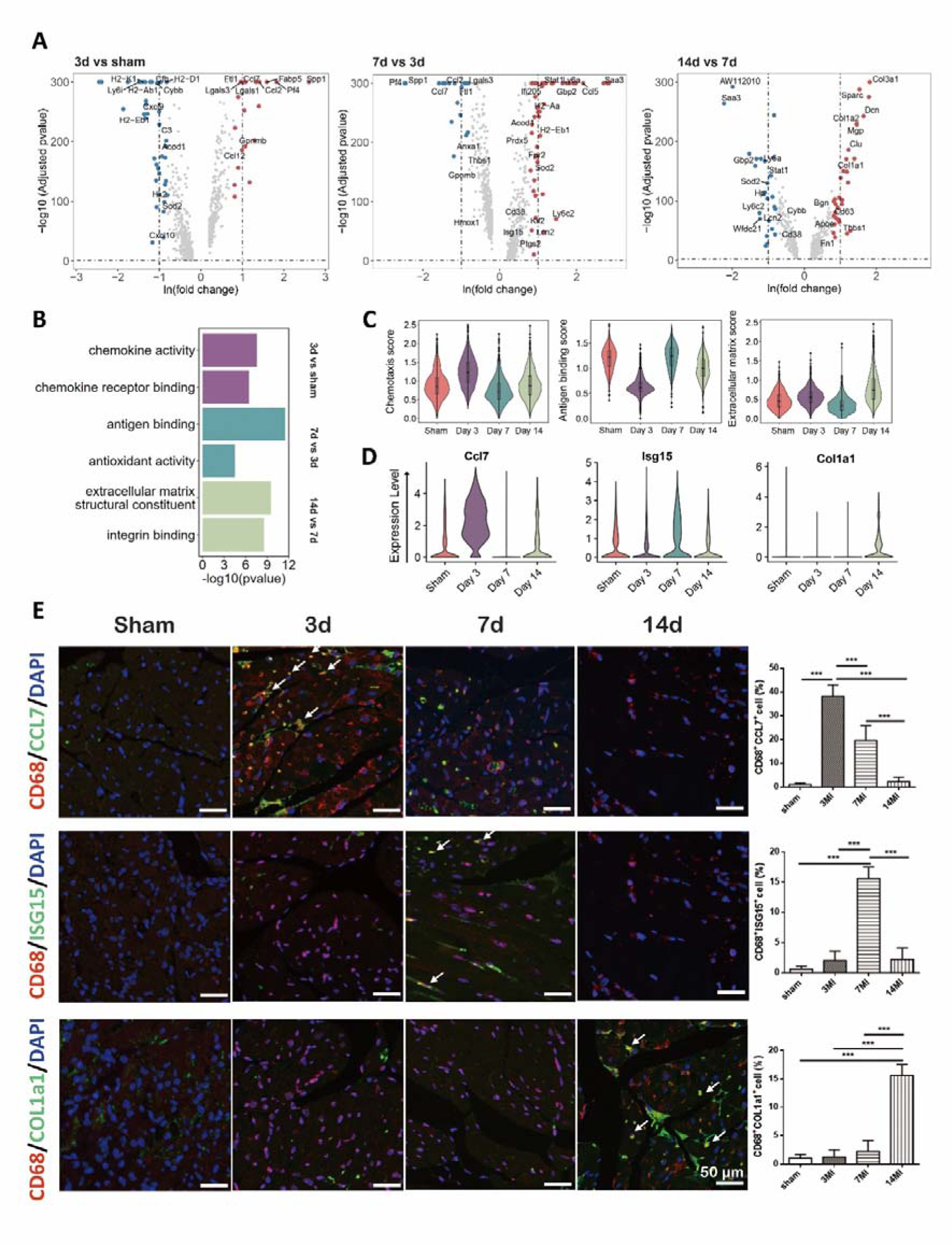
Time-specific molecular expression pattern of macrophage in MI progression. A. Volcano plot of the fold-change in differential gene expression in 3d MI and sham (left), 7d MI and 3d MI (middle) as well as 14d MI and 7d MI (right) from all cells in the Mø/monocyte subpopulation as shown in Fig. 1C. *P* values were calculated by Wilcoxon rank sum test and *P* adjusted values were corrected by Bonferroni analysis. B. GO enrichment terms of differentially expressed genes in 3d MI and sham, 7d MI and 3d MI as well as 14d MI and 7d. Detailed information is shown in **Figure S7A–C**. C. Temporal functional scores (chemotaxis (left), antigen binding (middle), extracellular matrix (right)) of all macrophages and monocytes. D. Violin plot of signature gene expression at each time point. E. Immunostaining images showing CCL7 (green), ISG15 (green), COL1a1 (green), and CD68 (red) positive cells in the heart. Nucleus was counterstained with DAPI (blue). White arrows indicate the CCL7^+^CD68^+^, ISG15^+^CD68^+^, or COL1a1^+^CD68^+^ cells in the heart. The percentages of CCL7^+^CD68^+^, ISG15^+^CD68^+^ or COL1a1^+^CD68^+^ cells were determined from images using ImageJ software.

### 2.5 Neutrophil subpopulations and their dynamic interactions with macrophages

Neutrophils are the primary cell type in the innate immune system. They infiltrate the heart and actively regulate postinfarction inflammation, profoundly affecting the outcomes of cardiac repair.^[7]^ Therefore, we further analyzed their heterogeneity and plasticity during MI progression. Three subsets with distinct expression profiles were obtained (**Figure 5A; Excel File S1**). Notably, these subsets were clearly distinguished over time (**Figure 5B; Figure S8A**). N1 specifically appeared on day 3 and expressed numerous chemokine genes (e.g., *Ccl2*, *Ccl7*, *Ccl9*, and *Ccl12*), triggering substantial extravasation of other immune cells in infarcted sites (**Figure S8B–C**). N2, another proinflammatory subset, exhibited minor expansion after infarction (**Figure 5B; Figure S8A**). N2 displayed strong up-regulation of interferon-stimulated genes and the granule protein-encoding gene *Mmp8*,^[24]^ which is also important for neutrophil-mediated host defense (**Figure S8B–C**). N3 was the most abundant subset on day 7 and differed by the expression of many genes corresponding to the apoptotic signaling pathway, such as *Sod2* and *Mif* (**Figure 5B; Figure S8A–C**). Using Monocle, we identified a possible aging trajectory for all neutrophils (**Figure 5C; Figure S8D**), with a progressive increase in the expression of age-related genes (*Icam1* and *Cd24a*) and reciprocal loss of the expression of *Cxcr2* and *Sell*^[7, 24]^ (**Figure 5D; Figure S8E**). In addition, N3 appeared at the end of trajectory and showed the highest aging and apoptotic scores (**Figure 5E**), suggesting that it was the most aged cluster.

**Figure 5.**
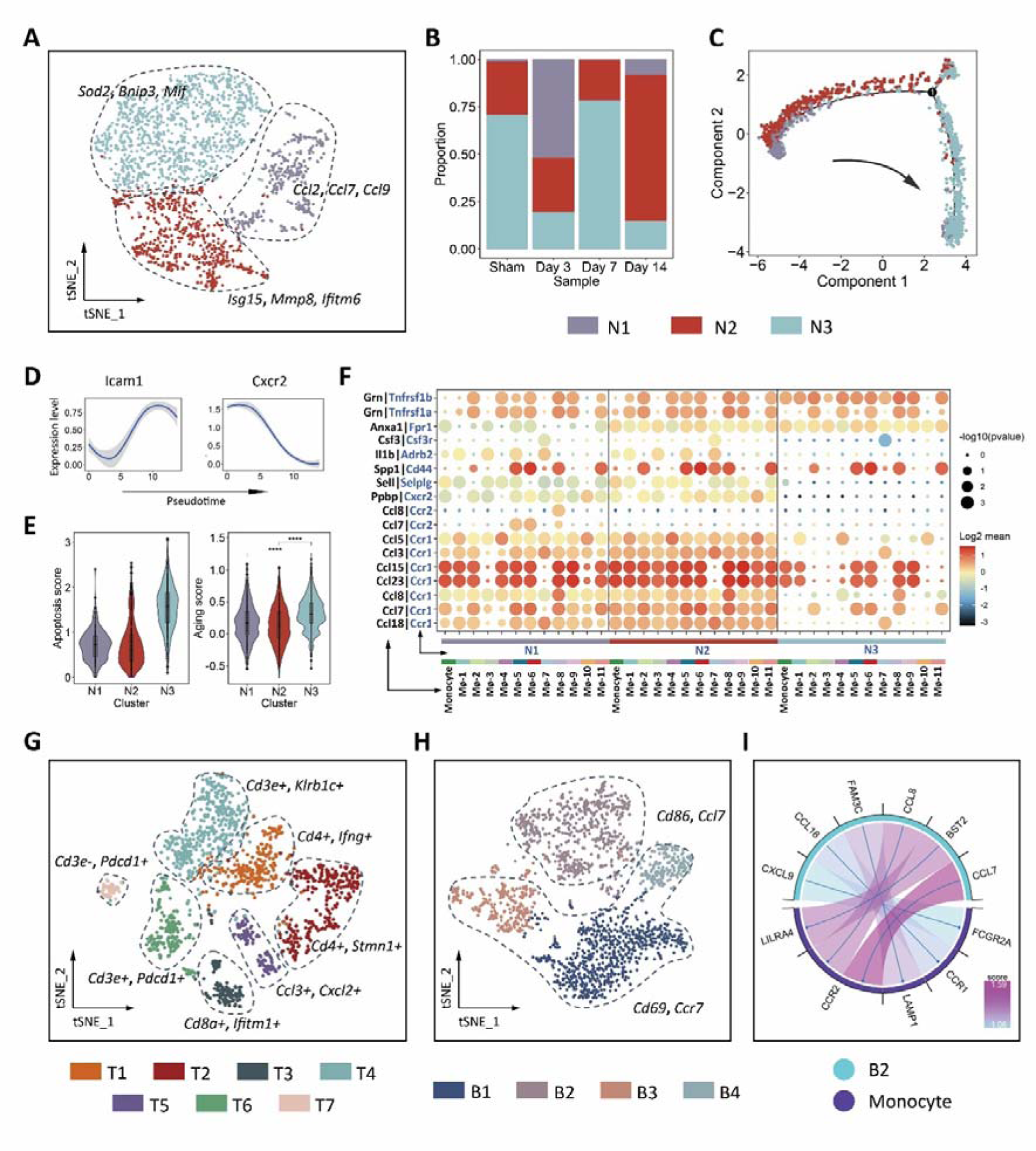
Temporal diversity and dynamics of neutrophils and lymphocytes. A. t-SNE plot of 1954 neutrophils showing three subsets merged from four samples in the MI dataset. All cells were obtained from the neutrophil subpopulation in Fig. 1C and reclustered to increase resolution. Feature genes in each subset are depicted in the plot. B. Fraction of each neutrophil subset relative to all cardiac neutrophils during MI progression. C. Pseudotime analysis revealed the age-related trajectory within three subsets of neutrophils. D. Expression of age-related genes was plotted along the pseudotime. E. Violin plot of apoptosis scores (left) and aging score defined as weighted average z-scores of age-related genes (right) for the three neutrophil subsets. Significance was determined by Wilcoxon test. NS, not significant (*P* > 0.05); *** *P≤* 0.001; **** *P* ≤ 0.0001. F. Overview of selected ligand–receptor interactions between Mø/monocyte and neutrophil; *P* values indicated by circle size, scale on the right. The means of the average expression level of interacting molecule 1 in cluster 1 and interacting molecule 2 in cluster 2 are indicated by color. t-SNE plot of 1445 T cells showing seven subsets (G) and 1866 B cells showing four subsets (H) merged from four samples in the MI dataset. All cells were obtained from T and B cell subpopulation in Fig. 1C and reclustered to increase resolution. Feature genes in each subset are depicted in the plot. I. Cell-cell interaction from B cell to monocyte.

We then used the CellPhoneDB algorithm to explore the crosstalk between neutrophils and macrophages (**Figure 5F**). As expected, macrophages showed increased interactions with N1 and N2 through ligand-receptor pairs associated with cell adhesion and chemotaxis, whereas N3 displayed reduced mobility. We also observed that macrophages secreted several anti-inflammatory ligands (*Grn*^[25]^ and *Anxa1*^[26]^), interleukin-1β (*IL-1*β), and colony-stimulating factor 3 (*Csf3*), which is important for extending the lifespan of neutrophils.^[27]^

Overall, the most abundant subset N3 that appeared on day 7 contributed to conversion from the inflammation phase to the repair phase after MI, given that dying neutrophils have the potential to limit postinfarction inflammation.^[2]^

### 2.6 Subpopulations of cardiac lymphocytes and their roles in MI

T- and B-cells are major components of myocardial lymphocytes. They coordinate adaptive immune responses, which are also vital for regulating ischemic injury.^[9]^ However, detailed dynamics in these immune cell types are less well-understood during MI progression. Therefore, we performed sub-clustering of these cells and generated t-SNE plots for visualization.

T cell populations in addition to Cd4/Cd8 T cells and NKT cells included several minor populations with distinct gene signatures: T5 (*Ccl3^+^ Cxcl2^+^*), T6 (*Cd3e^+^ Pdcd1^+^*), and T7 (*Cd3e^-^ Pdcd1^+^*) (**Figure 5G; Figure S9A–B, and Excel File S1**). Cd8 T cells were predominant at 3 days after MI (**Figure S9C**), indicating that these cells promote acute inflammation, leading to severe tissue damage.^[28]^ T4 was identified as NKT cells based on the expression number of markers (e.g., *Klrd1*, *Klre1*, and *Cd244*) as well as Cd3 (**Figure S9A–B**) and expanded dramatically by day 7 (**Figure S9C**). Cd4 T cells consisted of two distinct sets: T1 included typical active T cells expressing activation markers (e.g., *Cd4*, *Ifng*, *Cd28*, and *Cxcr3*) and enriched in T helper pathways in Ingenuity Pathway Analysis, and T2 included cycling Cd4 T cells, characterized by up-regulation cell cycle-related factors (**Figure S9D–E**). T6 and T7 expressed high levels of *Pdcd1* (**Figure S9E**), which encodes the T cell exhaustion marker PD-1 and is generally considered as an anti-tumor target. However, increasing evidence shows that upregulation of this gene limits excessive immunopathology and has a cardiac protective function.^[29]^

Finally, we clustered B cells into four subpopulations (**Figure 5H; Excel File S1**). B2 existed at all time points but showed minor expansion after infarction (**Figure S10A**). This cluster exhibited differential expression of the activation marker *Cd86* and chemotactic gene *Ccl7* (**Figure S10B**). B cell *Ccl7* mediates the recruitment of circulating monocytes, which enhances tissue injury and leads to the deterioration of myocardial function.^[20c]^ This type of cellular crosstalk was confirmed by our cell–cell interaction analysis (**Figure 5I**). The remaining three clusters increased and peaked at around 7 days after MI (**Figure S10A**) with high levels of the B cell activation markers *Cd69* and *Ccr7* (**Figure S10B**), a chemoattractant receptor, which may enhance B cell antigen presentation and migration to T cell zones.^[30]^ These findings suggest that an interaction occurs between T and B cells.

#### 2.7 Suppressing early infiltrated Mø-5 and Mø-6 attenuates pathological MI progression

Day 3 is characterized as the acute inflammatory stage and may be vital for disease modulation. Given that macrophages expanded dramatically at this time point (**Figure 1E**), we focused on this population to identify critical subsets that allow time- and cell type-specific intervention in the pathogenesis of MI. The disease contributions of different macrophage subsets were first determined by considering the alterations in both the cell number and gene expression with the contribution score, using scRNA-seq data from the “sham” and “3d MI” groups (detailed method is described in the **Experimental Methods**). As shown in **Figure 6A**, infiltrated Mø-5 and Mø-6 exhibited the highest contribution score among all macrophage subsets, indicating that they are crucial for aggravating disease progression. To explore whether targeting these two subsets can alleviate ischemia impairment and prevent further injuries, we applied tanshinone IIA, a well-known natural cardioprotective agent against MI that was isolated from the traditional Chinese medicinal plant *Salvia miltiorrhiza*,^[31]^ as a proof of concept (**Figure S1A**). Namely, we integrated our original MI dataset with the scRNA-seq data of tanshinone IIA-treated murine hearts that were parallelly acquired during MI experiments (**Figure 6B**). Mapping of the resulting integrated dataset to the annotated MI dataset was performed using reference-based supervised algorithms.^[32]^ Similar immune cell populations and their precise subsets that were matched to the MI dataset were identified (**Figure S11A–E**). Notably, administration of tanshinone IIA substantially reduced the Mø/monocyte population on day 3 (**Figure 6C**).

**Figure 6.**
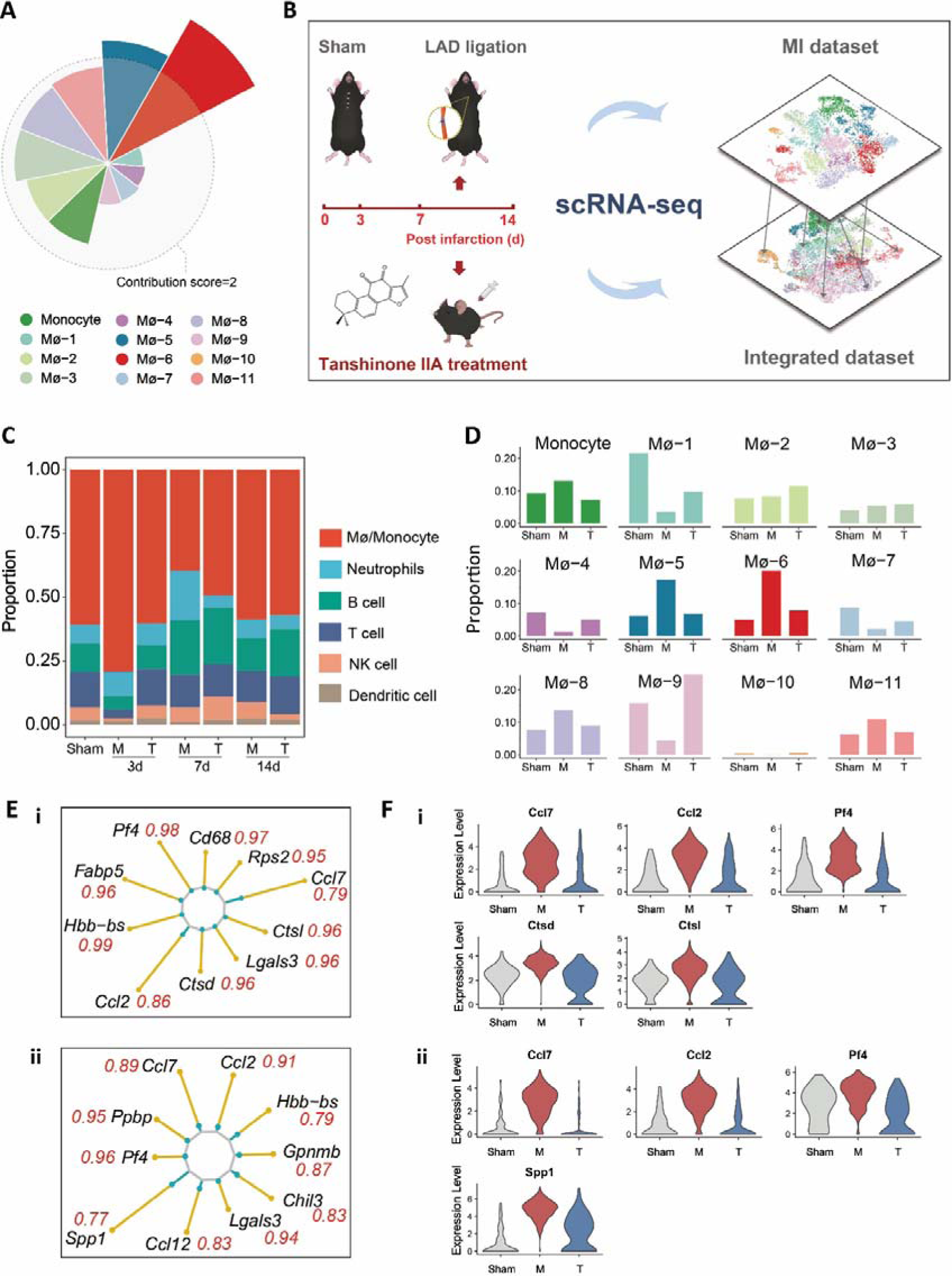
Tanshinone IIA treatment validates the effectiveness of early infiltrated macrophage subsets suppression in attenuating pathological MI progression. A. Disease contributions of distinct macrophage subsets. The radius is proportional to the contribution score. Detailed information is described in the **Experimental Methods**. B. Schematic diagram of the experimental design and data integration. 10X chromium scRNA-seq was performed on *Cd45*^+^ immune cells isolated from the parallel tanshinone IIA-treated hearts of C57/BL mice at 3, 7, and 14 days after left anterior descending artery ligation. Five mice at each time point were processed simultaneously. scRNA-seq data of Tan IIA groups was pre-processed using the same pipeline and integrated with the MI dataset. C. Fraction of immune cell populations relative to all cardiac immune cells as a function of disease progression and tanshinone IIA treatment. D. Alterations in the proportion of monocyte and each macrophage subsets after tanshinone IIA treatment on day 3. E. Top 10 genes suppressed by tanshinone IIA in Mø-5 (i) and Mø-6 (ii). The length of yellow and blue lines indicates the gene expression level change in the “3d MI” or “3d Tan IIA” group compared to that in the sham group, respectively. The recovery regulation ratio (*Rr*) of each gene is depicted in red. F. Violin plot of signature gene expression changes after tanshinone IIA treatment on day 3 in Mø-5 (i) and Mø-6 (ii).

More importantly, Mø-5 and Mø-6 were the two predominant subsets for which the cell proportion was dramatically decreased by 60% after tanshinone IIA treatment (**Figure 6D**), indicating their potential in targeting macrophage subsets for intervention of MI. In-depth analysis of the top differentially expressed genes with the recovery regulation ratio (Rr) revealed that the expression levels of several chemokines (e.g., *Ccl7*, *Ccl2*, and *Pf4*) were significantly decreased in both the Mø-5 and Mø-6 subsets after tanshinone IIA treatment compared to in the MI group (**Figure 6E–F**). Several cathepsins (e.g., *Ctsd* and *Ctsl*) associated with inflammation in Mø-5 showed lower expression in the tanshinone IIA-treated group compared to in the MI group, and the expression level of the multifunctional cytokine *Spp1* which is essential for resolving inflammation and initialing the scar deposition^[33]^ was also reduced in Mø-6 after tanshinone IIA treatment (**Figure 6F**). These results indicate that both the cell proportion and functional genes of Mø-5 and Mø-6 are targeted to alleviate MI.

Additionally, to investigate whether suppressing early infiltrated pro-inflammatory Mø-5 and Mø-6 influences the subsequent outcomes of disease progression, we also examined detailed leukocyte alterations after tanshinone IIA treatment at 7 and 14 days after MI. Interestingly, we detected a large number of neutrophils that disappeared on day 7 (**Figure 6C**). Deeper analysis revealed that most of these cells were originally from N3, the previous defined apoptotic subset (**Figure S11F–G**). On day 14, although the proportion of Mø-10 associated with collagen formation was increased, the expression levels of collagen-related genes in this set were suppressed by tanshinone IIA (**Figure S11H–I**), consistent with the result of Masson’s trichrome staining (**Figure S2A**). These results demonstrate the effectiveness of suppressing early infiltrated macrophage subsets (Mø-5 and Mø-6) in attenuating pathological MI progression, indicating the importance of cell type- and time-specific intervention.

## 3. Discussion

Although several studies have provided scRNA-seq data on MI and insight into the complex cardiac immunity,^[6-7, 11a, 13, 29b]^ systematic analysis of post-MI immune cell dynamics is required. Therefore, we analyzed all *Cd45*^+^ cells present in the hearts of mice with left anterior descending artery ligation at different time points (0, 3, 7, and 14 days) by scRNA-seq. Our findings provide information on the immune landscape at infarcted sites for a wide range of cell populations over a broad time span.

Macrophages are the most abundant cardiac immune cells and play a central role in multiple pathophysiological processes of MI.^[5b]^ Increasing evidence has highlighted the heterogeneity of macrophages resulting from their different origins and functions.^[5a, 10]^ Nevertheless, an arbitrary M1–M2 dichotomy has been commonly used to group these cells into the detrimental inflammatory population and opposite protective anti-inflammatory population, which may be associated with tissue repair and wound healing.^[34]^ Even in some recent studies using scRNA-seq, simplified classification focused only on inflammatory-related marker genes has been performed.^[8, 29b, 35]^ However, in this study, we defined 11 macrophage subsets in both homeostatic and disease states, each with a distinct function according to their unique gene expression profiles and GO enrichment analyses (**Figure 2B; Excel File S2**).

We analyzed macrophage dynamics during MI progression and revealed critical subsets and functionally important genes as potential therapeutic targets against MI. Notably, Mø-5 and Mø-6 were defined as the two major postinfarction subsets at 3 days after MI, and they exhibited strong expression of chemokine genes (*Ccl7*, *Ccl2*, and *Pf4*). *Ccl7* was reported to be expressed in tissues showing active inflammatory reactions and is strongly associated with *Ly6c*^+^ monocytes infiltration.^[36]^ In patients with acute MI, high concentrations of circulating CCL7 have been observed and are considered to increase the risk of death or recurrent MI.^[20c]^ Similarly, an elevated CXCL4 (CXC chemokine ligand-4; *Pf4*) plasma level in patients with acute coronary syndrome was observed.^[37]^ Recent studies showed that CXCL4 is closely related to increased matrix metalloproteinases, which account for cardiac remodeling.^[20b, 38]^ However, these highly amplified macrophages (Mø-5 and Mø-6) and their expressed inflammatory factors were markedly suppressed by tanshinone IIA treatment, indicating that both the macrophage subsets and functionally important genes are potential therapeutic targets in MI progression. We also found that Mø-6 expressed a high level of *Spp1*, which is crucial for inflammatory restraint and collagen deposition.^[33]^ Down-regulated expression of this gene in Mø-6 was observed after tanshinone IIA treatment (**Figure 6F**). A recent study reported that nearly all *Spp1* is produced by galectin-3^hi^CD206^+^ macrophages after myocardial injury.^[21]^ However, our data demonstrated that *Spp1* is expressed in not only CD206^+^ macrophages, but also pro-inflammatory subsets, suggesting that classic “M1” macrophages can promote a reparative fibrotic response. As the major postinfarction subsets Mø-5 and Mø-6 were crucial for disease progression, we also explored the developmental relationships between these cells and monocytes. Mø-5 and Mø-6 were mainly derived from monocytic progenitors except for some fraction of Mø-6. This fraction expressed lower *Ccr2* and appeared more distant from the monocyte in the developmental trajectory (**Figure 3B**), suggesting that they can arise from tissue-resident macrophages. However, *Ccr2* was highly expressed in some resident subsets on day 3 (**Figure S6D**), demonstrating that they may also be replenished from circulating monocytes after injury. A recent study revealed that monocyte-derived macrophages can gain multiple cell fates, even those that are difficult to distinguish from resident macrophages.^[6]^ Therefore, further studies of these differentiation processes is required to determine the origins of postinfarction macrophages, which is important for developing precise interventions for MI.

At 7 days, we found that macrophage subsets with inflammatory features were still prominent (**Figure 3B**). This may be because these sets highly express *MHC* genes and are essential for inducing adaptive immune responses, consistent with a previous study.^[8]^ The apparent expansion of lymphocytes at the same time was observed in our dataset (**Figure 1F**), and some cells showed a cardiac protective function. Additionally, we detected a large apoptotic subset of neutrophils on day 7 after MI, which was nearly lost by day 14 (**Figure 5B; Figure S8A**). This type of clearance is an important cellular mechanism for attenuating inflammation and may activate anti-inflammatory programs, contributing to the conversion to a reparative stage in the ischemic heart. As a result, we observed that the proportion of macrophage subsets involved in tissue repair was increased in the final stage. Particularly, Mø-10 expressing many collagen-related genes specifically appeared at this time point (**Figure S6B**). The cell number of this subset increased after tanshinone IIA treatment; however, the average expression level of some profibrotic genes was suppressed (**Figure S11G–H**). Therefore, further studies are required to determine beneficial or adverse contribution of this subset to cardiac remodeling so that effective therapies can be developed.

Our study provides insight into the systematic post-infarction dynamics of cardiac immune cells in the progression of MI. Mø/monocytes, neutrophils, and lymphocytes dominated at different stages and showed distinct roles. Early infiltrated macrophages and their expressed inflammatory factors played an important role in the pathogenesis of MI. Neutrophil apoptosis and lymphocyte expansion facilitated the conversion from the inflammatory into the reparative phase on day 7. A profibrotic macrophage subset specifically appeared at 14 days, which can influence the outcomes of cardiac remodeling. Treatment with the active compound suggested that cell type- and time-specific interventions have potent clinical applications in MI.

## 4. Experimental Methods

### 4.1 Animal experiment

All animal studies were approved by the Laboratory Animal Ethics committee of Tianjin University of Traditional Chinese Medicine (Permit Number; TCM LAEC2014004). All animal experiments were performed on male 7–8-week-old C57/BL mice (weighing 20–24 g) purchased from Vital River Laboratory Animal Technology Co., Ltd. (Beijing, China). Mice were allocated to seven groups at random: (1) Sham group (Control, n = 5); (2) 3d MI group (n = 5); (3) 3d Tan IIA group (n = 5); (4) 7d MI group (n = 5); (5) 7d Tan IIA group (n = 5); (6) 14d MI group (n = 5); and (7) 14d Tan IIA group (n = 5). To induce MI, permanent ligation of the left anterior descending artery was applied. For the Tan IIA groups, the mice were intragastrically administered with 20 mg/kg tanshinone IIA (Solarbio, ST8020) daily. Mice from different groups were examined by echocardiography and sacrificed at 0, 3, 7 and 14 days after surgery. Excised hearts were processed for immune cell isolation (n = 5, per group).

### 4.2 Left anterior descending coronary artery (LAD) ligation

MI was induced by permanent ligation of the LAD. Briefly, mice were anesthetized by intraperitoneal injection of 15 mg/mL tribromoethanol (0.01 mL/g) (Sigma-Aldrich, T48402) and placed in a supine position on a temperature control pad. Then, the animals were intubated and ventilated with a 16-gauge intravenous catheter. Left-sided thoracotomy was performed by small incision between the third and fourth intercostal spaces. The incision was expanded by a blunt-ended retractor to avoid the retraction of the lungs. After exposing the heart, a cotton bud was gently pressed on the artery a little below the ligation site of LAD (8 mm away from the origin). A 5-0 silk ligature was passed underneath the LAD and tied with three knots using a tapered atraumatic needle. Visible blanching and cyanosis of the anterior wall of the left ventricle and swelling of the left atrium were indicative of successful ligation. Ribs and muscles were closed using 6.0 vicryl dissolvable suture leaving a small gap to aspirate air left in the chest cavity.

### 4.3 Echocardiography

To assess cardiac function, echocardiography was performed on anesthetized mice (full anesthetic; 1 % oxygen plus 5 % isoflurane anesthetic in the anesthetic box) from MI or Tan IIA groups at different time points (0, 1, 3, 7, and 14 days after the surgery), using a Vevo2100 system (Visualsonics, US) fixed with the MS-400 probe. The M-type ultrasound image was obtained, the long shaft measuring package (PLAX) was selected to measure the shrinkage and diastolic period of the mice. The interventricular septum thickness (IVS), the left ventricular internal diameter (LVID), the left ventricular ejection fraction (EF), fractional shortening (FS), left ventricular mass, and left ventricular volume were measured. 3 independently acquired images of each mouse were obtained to calculate the mean value. After the surgical group ultrasound, the EF value in the range of 38 to 50 % was considered to be successful.

### 4.4 Flow cytometry

Mice in shame and MI groups were anesthetized by intraperitoneal injection of 15 mg/mL tribromoethanol (0.01 mL/g) (Sigma-Aldrich, T48402) on day 0, 3, 7, and 14. Following taking the abdominal aortic blood, 5 mL pre-cooling PBS was used to perfuse. Then the hearts were cut up into small pieces in a petri dish filled with pre-matched digestive fluid and incubated in a shaker at 37 °C, 100 rpm for 15 min. Cell suspensions were filtered by 75 μm mesh twice, added 1 mL complete medium, and centrifuged at 1500 rpm for 5 min. After discarding the supernatant, the cell pellets were suspended in the mixture of 1 mL erythrocyte lysate and 4 mL of PBS. Then centrifuge at 1500 rpm for 5 min and discard the supernatant. Cell pellets were resuspended in 500 μL PBS and incubated with antibodies against murine CD11b (BD Biosciences, 553310), F4/80 (BD Biosciences, 565787), and Ly6C (Abcam, ab24973) in 4 °C for 30 min. After adding 2 mL PBS and washing twice. 500 μL 1% paraformaldehyde was added to fix the cell in 4 °C. Flow cytometry analysis was performed on the second day.

### 4.5 Immunofluorescence

Heart tissues were fixed in 4 % paraformaldehyde and embedded with paraffin. Sections (5 μm thickness) were cut in to microscopic slides. Paraffin-embedded heart samples were dewaxed and rehydrated in xylene and ethyl alcohol, followed by incubation in 0.3% methanol to endogenous peroxidase. Then it was rehydrated and permeabilized in PBS with 0.1% (v/v) triton X-100/0.25% BSA. Antigen retrieval was performed by boiling the slides in a microwave oven with an antigen retrieval solution for 20 min. Next, sections were incubated overnight at 4 °C with mouse anti CD68 (1□100) (Abcam, ab201340), rabbit anti CCR2 [(1□100) (Abcam, ab216863), recombinant anti CCL7 (1□0) (Abcam, ab228979), ISG15 recombinant rabbit monoclonal antibody (1□00) (Invitrogen, 703132), and collagen I (1□100) (Affinity Biosciences, AF7001) primary antibodies at the dilution indicated. 488 or 594 conjugated anti-rabbit secondary antibody was applied following primary antibody incubation. The sections were counterstained with DAPI (Beyotime, P0131) and examined under a fluorescence microscope (Leica).

### 4.6 Histopathology

To evaluate the morphological changes of the heart and the extent of cardiac fibrosis, murine hearts from different groups were harvested, washed in PBS and fixed in 4% paraformaldehyde overnight and then embedded in paraffin solution. Each heart was cut into 4-μm-thick sections and stained with hematoxylin and eosin staining (HE), Masson’strichrome-staining.

### 4.7 Tissue collection and *Cd45*^+^ cells isolation

Infarcted areas (infarct and border zone region) from the MI or administrated murine heats and the corresponding region of control hearts were collected and minced into fine pieces. Tissues were enzymatically digested with a multi-tissue dissociation kit (Miltenyi Biotec, 130-110-203), according to the manufacturer’s instructions. After dissociation, cell suspensions were passed through a 30 μm Cell Strainer, followed by red blood cells lysis (Solarbio, R1010). Dead cells and debris were removed by using the dead cell removal kit (Miltenyi Biotec, 130-090-101). Then single cell suspensions were incubated with CD45 MicroBeads, mouse (Miltenyi Biotec, 130-052-301) for 10 minutes at 4 °C, and isolated by a MACS Separator (Miltenyi Biotec).

### 4.8 Single cell library preparation and sequencing

After washing and resuspending in an appropriate volume of cold PBS, purified *Cd45*^+^ cells were processed in the Chromium single cell gene expression platform (10x Genomics). In brief, Single Cell 3′ Reagent Kit v2 (10X Genomics) was used to generate droplets encapsulating single cell and barcoded bead. Following lysis, reverse transcription was performed in a thermal cycler (ProFlex™ PCR). cDNA was purified with Dynabeads (provided by 10x) and amplified for 10 or 14 cycles (ProFlex™ PCR). The amplified-cDNA was fragmented, ligated with adapter and sample index, and selected with SPRI beads (Beckman). The constructed library was sequenced on the NovaSeq 6000 Illumina sequencing platform.

### 4.9 Quantification, quality control, and cluster analysis of single-cell expression

We aligned and quantified the 10X raw sequencing data using the CellRanger (10X Genomics, version 3.0.2) suite. The reads were mapped by the CellRanger count module with a mouse genome reference (mm10). **Table S1** showed the global mapping report of the MI dataset, including Estimated Number of Cells, Mean Reads per Cell and Mean Genes per Cell for each experimental time point.

Raw gene expression matrices were filtered by Seurat R package^[39]^ (version 3.1.5) and the cells that expressed less than 200 or more than 5000 unique genes or more than 20 of reads mapping to mitochondria were filtered out. The global-scaling method was used to normalize the gene expression of the remaining 17,932 cells with a default scale factor.

The FindVariableGenes function in Seurat was used to select high variable genes for integrating samples from the MI dataset with the function IntegrateData. The normalized expression was scaled through the function of ScaleData to remove unwanted sources of variation. After principal component analysis (PCA), the most significant 20 principal components (PCs) were used for tSNE dimensionality reduction. Cells were clustered by the FindClusters function, and each cluster was annotated by the expression of known marker genes. To focus on immune cells, we filtered out the *Cd45*^-^ cells and performed downstream analysis on the remaining 16,380 cells (**Figure S3B**).

### 4.10 Pseudotime analysis

The normalized gene expression matrix was used as an input to Monocle2.^[40]^ A new object of Monocle2 was created by the newCellDataSet function and gene expressions were updated by the dispersionTable function. The genes with a mean expression lower than 0.001 and the genes detected in less than 10 cells were filtered. After dimension reduction, the cell trajectory was inferred according to the 1000 most variable genes selected by FindVariableFeatures function in Seurat with the method of “vst”.

Similar to the methods used in Monocle2. the normalized gene expression matrix was used as an input to Monocle3^[41]^ to infer the potential lineage differentiation trajectory. A new object of Monocle3 was created by the new_cell_data_set function and processed by the preprocess_cds function with default parameters. We reduced dimensionality of data using the reduce_dimension function (umap.min_dist = 0.5, umap.n_neighbors = 35) before learning the trajectory and order of cells.

### 4.11 Cell-cell communications analysis

CellPhoneDB is a tool based on gene expression and curated knowledge of communication information, such as receptors, ligands and their interactions, from known databases. The genes that expressed in less than 10 cells were filtered out and mouse genes were converted to human paralogues by R package biomaRt^[42]^ (version 2.42.1) with the GRCm38/GRCh38 genome reference. The expression matrix was imported in CellphoneDB^[43]^ (version 2.1.4) and performed analysis. In the results of CellPhoneDB, we filtered out the ligand-receptor pairs that had no efficient mean interacting values.

### 4.12 Gene Ontology enrichment analysis

The FindAllMarkers function was used to find out the differential genes of each cell subset by comparing the cell subset with other cells. For downstream analysis, differential genes were filtered by following settings (only.pos = TRUE, min.pct = 0.2, logfc.threshold = 0.2) and a list of differential genes at an adjusted *p* value (Wilcoxon test) < 0.05 were retained.

We selected the top 100 genes in fold change to Perform Gene Ontology(GO) enrichment analysis using clusterProfiler R package.^[44]^ The results of the enrichment analysis were selected based on the statistical threshold (qvalueCutoff = 0.05) and the results belonging to Biological Process (BP) were reserved.

### 4.13 Ingenuity pathway analysis

The differential genes list of each cluster was calculated with the FindAllMarkers function (only.pos = FALSE, min.pct = 0.2, logfc.threshold = 0.2). The differential genes at an adjusted p value cut-off of 0.05 (Wilcoxon test) provided with p-value and fold change were imported into IPA (QIAGEN Inc., https://www.qiagenbioinformatics.com/products/ingenuity-pathway-analysis) for ingenuity pathway analysis.

### 4.14 Scoring of biological processes

The scores of individual cells were defined as the average normalized expressions of gene signatures representing distinct biological functions. Functional signatures were collected from the Gene Ontology database and they are all differential genes at an adjusted p value cut-off of 0.05 using Wilcoxon test. For “ROS metabolic process” and “Response to oxidative stress”, the score was calculated according to the genes enriched in corresponding pathway

### 4.15 Integrating scRNA-seq data of Tan IIA groups with MI dataset and inferring the cell type

We pre-processed scRNA-seq data of Tan IIA groups via the CellRanger (10X Genomics, version 3.0.2) suite. **Table S3** showed the mapping results. After using the same pipeline implemented in the Seurat R package, we created an integrated dataset that combined the new data with the original MI dataset. We then performed the unsupervised clustering and inferred the cell type using SingleR^[32a]^ (version 1.0.6). Briefly, the original MI dataset with known cell types was used as the reference dataset. Each cell in the integrated dataset was performed annotation independently, according to their gene expression similarity to that in the MI dataset. The cell type annotation of each cluster in the integrated dataset was determined by the maximum assigned cell type.

As for Mø/monocyte population in the integrated dataset, we performed clustering and cell type annotation using ItClust,^[32b]^ which is an Iterative Transfer learning algorithm for scRNA-seq Clustering. Gene expression signatures in the well-labeled MI dataset were used to build a training neural network. Clustered cells with assigned cell type were returned from the target network after finishing fine-tuning.

### 4.16 Characterization of MI contribution in different macrophage subsets

We characterized the disease contribution of distinct macrophage subsets by considering both cell number and gene expression variations. We first determined the signature genes of each subset. Briefly, we performed the bulk differential gene expression analysis and selected the top 100 up-regulated genes from the “sham” to the “3d MI” group to characterize the alteration during the process. Besides, we analyzed the differentially expressed genes between “sham” and “3d MI” groups in each subset separately. Those which also belong to the top-100 bulk differentially expressed gene set was chosen as the signature genes for each set. Then, we defined the *FC_score_* that measures the quantity and expression level change of signature genes during the biological processes.

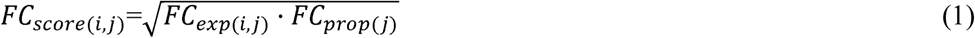

In which *FC_exp(i,j)_* refers to the expression fold change of the i^th^ signature gene in j^th^ cluster, *FC_score(i,j)_* is the proportion fold change of the j^th^ cluster. of the i^th^ signature gene in j^th^ cluster is defined as the square root of, *FC_exp(i,j)_* by *FC_exp(i,j)_*.

Finally, the MI contribution of distinct macrophage subsets was characterized by the average *FC_score_* of all signature genes in this cluster.

### 4.17 Calculation of the regulating score (RS) and the recovery regulation ratio (Rr)

We defined the regulating score (*RS*) of genes to characterize their expression level changes compared to that in the “sham” group.

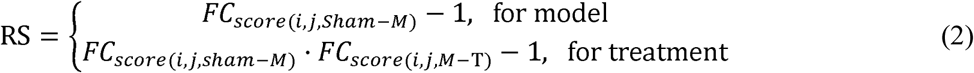

In which *FC_score(i,j,sham-M)_*, refers to the *FC_score(i,j)_*, of genes during the process from “Sham” to “3d MI” group. Likewise, *FC_(i,j,M-T)_*, refers to the *FC_score(i,j)_*, of genes during the process from “3d MI” to “3d Tan IIA” group.

We then defined the recovery regulation ratio (Rr) by:

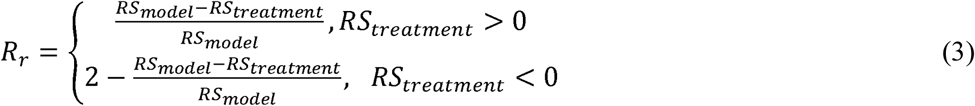

The closer Rr is to 1, the stronger regulation of tanshinone IIA.

## 5. Supporting Information

Supporting Information is available from the Wiley Online Library or from the author.

## 6. Acknowledgements

The authors gratefully acknowledge OE Biotech for assisting with scRNA-seq experiments. This work was supported by the National Natural Science Foundation of China [81973701 to X.F. and 81904054 to S.G.], the Natural Science Foundation of Zhejiang Province [LZ20H290002 to X.F.], the National Youth Top-notch Talent Support Program [W02070098 to X.F.], the National Key Subject of Drug Innovation [2019ZX09201005-007 to G.F.], and the Tianjin Science Foundation for Distinguished Young Scholars [17JCJQJC46200 to G.F.].

