## Supplemental Information for "Single-cell RNA Sequencing Reveals the Temporal Diversity and Dynamics of Cardiac Immunity after Myocardial Infarction"

**This PDF file includes:**

Figs. S1 to S11

Tables S1 to S3

**Other Supplementary Materials for this manuscript include the following:**

Excel File S1 to S3

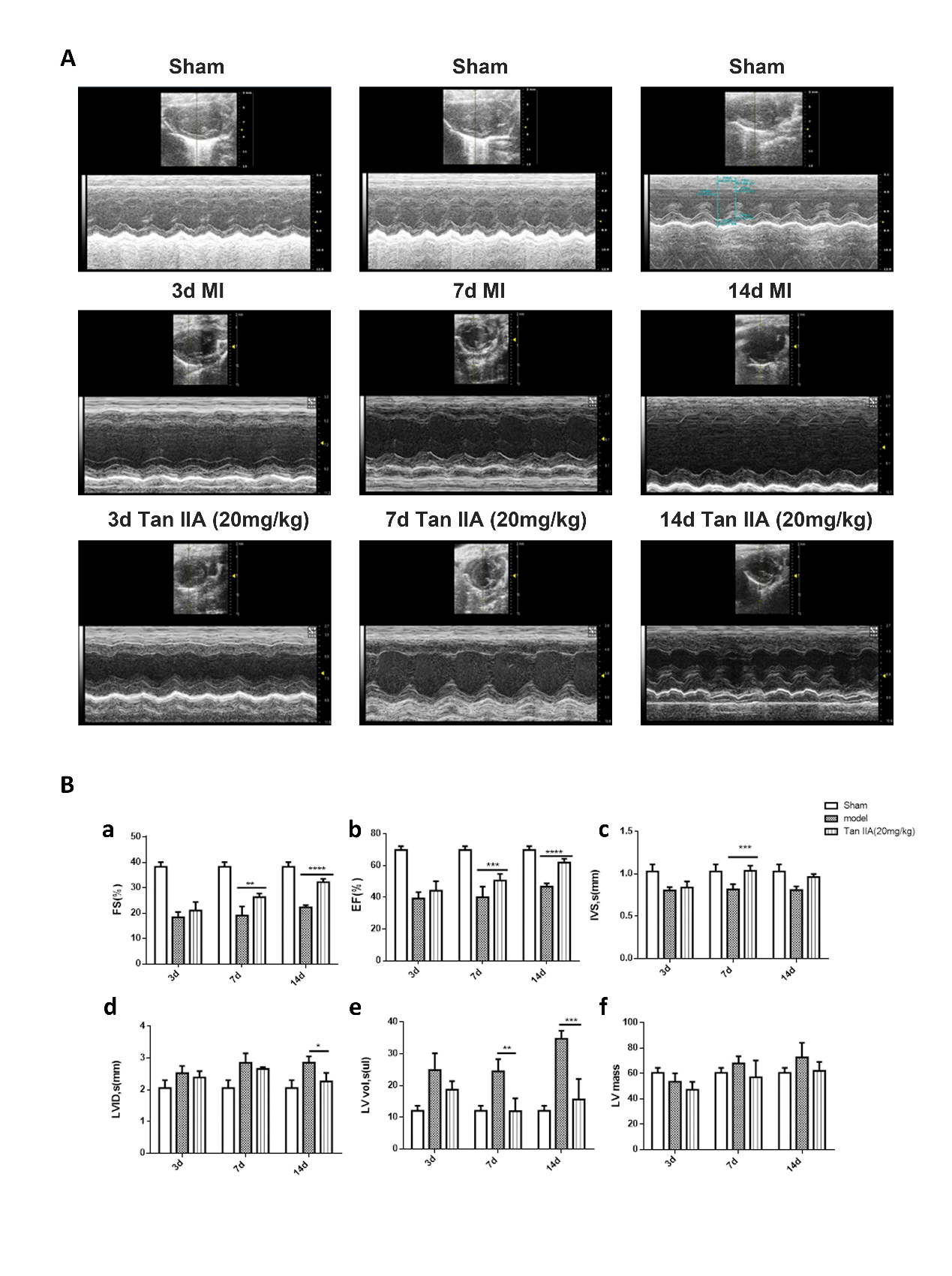

Fig. S1.

(A) Echocardiography assessment of cardiac functions in mice from MI and Tan IIA groups at 3, 7, and 14 days after the surgery. (B) Left ventricular fractional shortening (a), left ventricular ejection fraction (b), thickness of the left anterior ventricular wall (c), ventricular inner diameter (d), ventricular volume of the heart (e), and ventricular size and quality of the heart (f) in each group. Data expressed as mean ± SD, n = 5. * *P* < 0.05 vs model, ** *P* < 0.01 vs model, *** *P* < 0.001 vs model.

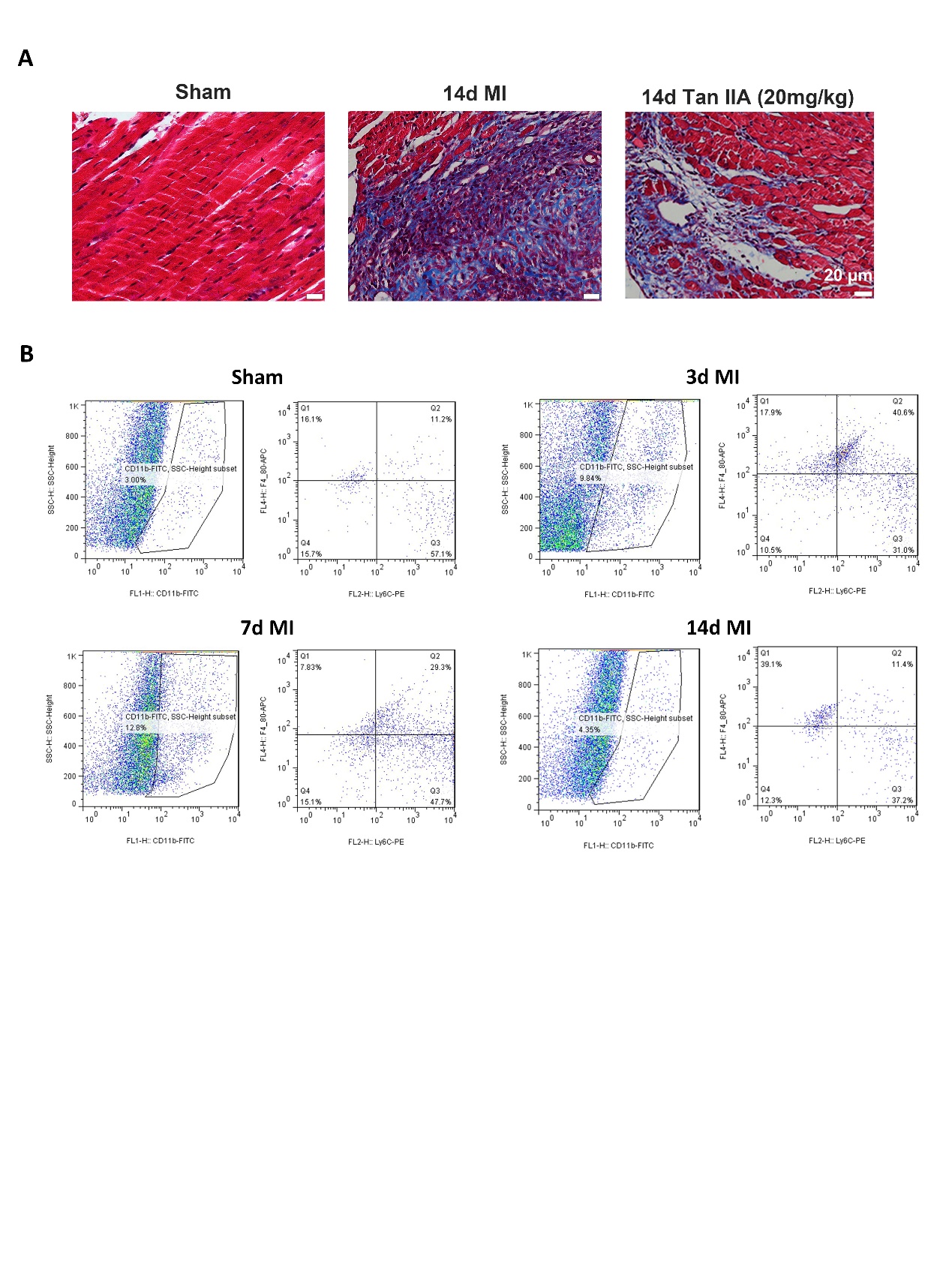

Fig. S2.

(A) Representative illustration of Masson’s trichrome-staining of infarcted mouse hearts in MI or Tan IIA group on day 14 and the control heart. (B) Flow cytometry analysis showed temporal changes of monocytes and macrophages at different days after MI.

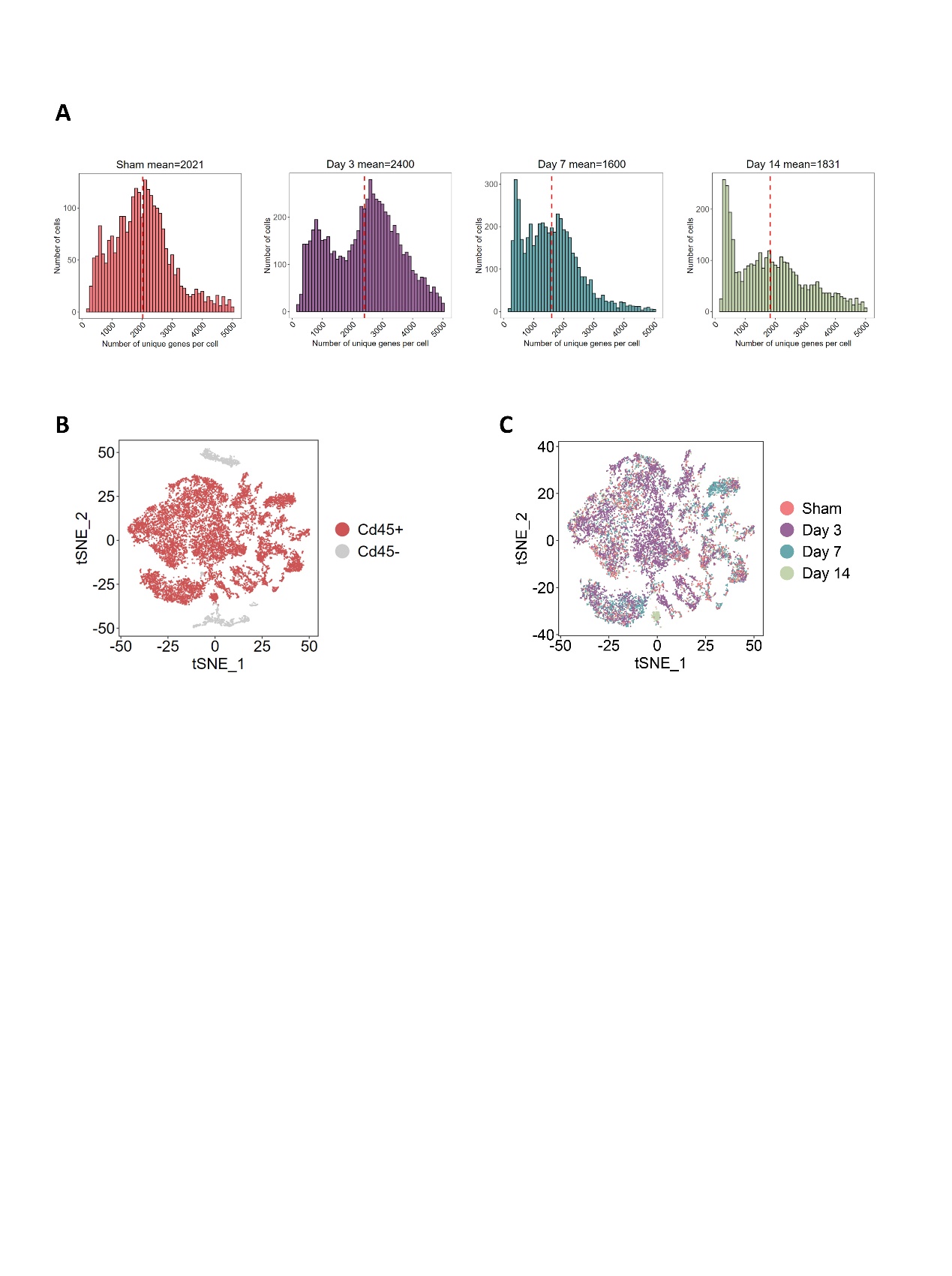

Fig. S3.

(A) Distributions of the number of genes detected per cell at different stages after MI. This dataset was quality trimmed and filtered. Red dashed lines indicate mean values. (B) tSNE plot showing the *Cd45*^+^ and *Cd45*^-^ cells from four time points. (C) tSNE plot of 16,380 *Cd45*^+^ cells visualized by different origins.

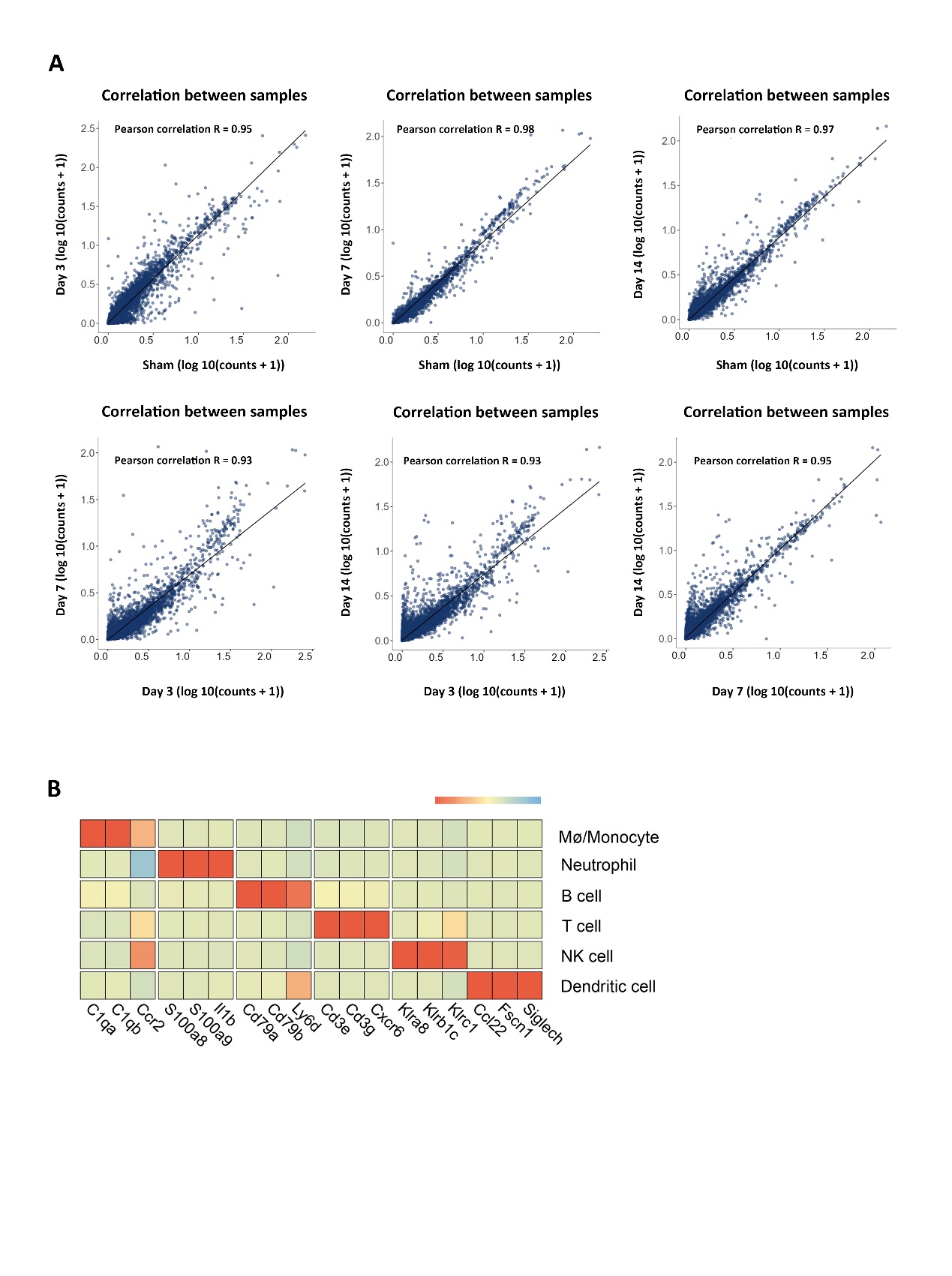

Fig. S4.

(A) Scatterplots showing Pearson correlation of the gene expression detected among four samples (i.e., Sham, 3d MI, 7d MI, and 14d MI). *P*-values are all less than 0.05. The analysis was conducted aggregating the cell information across corresponding samples. (B) Heatmap of normalized expression for selected marker genes in different immune subpopulations.

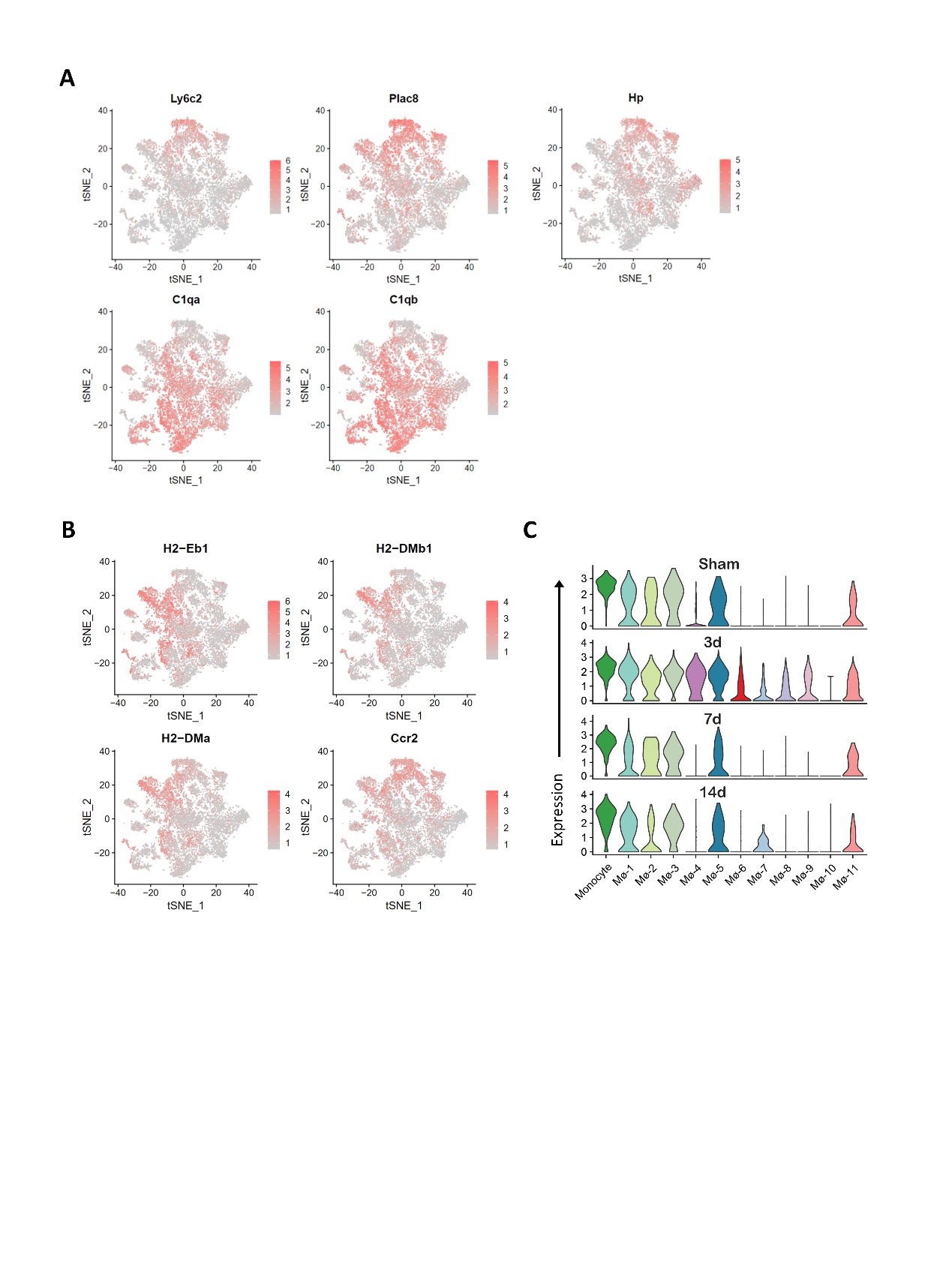

Fig. S5.

(A) Feature plot showing monocyte and macrophage marker gene expression gradients identified within each subset. (B) Feature plot of *MHC* genes (*H2-Eb1*, *H2-DMb*, and *H2-DMa*) and *Ccr2* in the Mø/monocyte population. (C) Violin plot showing dynamic expression levels of *Ccr2* within each subset.

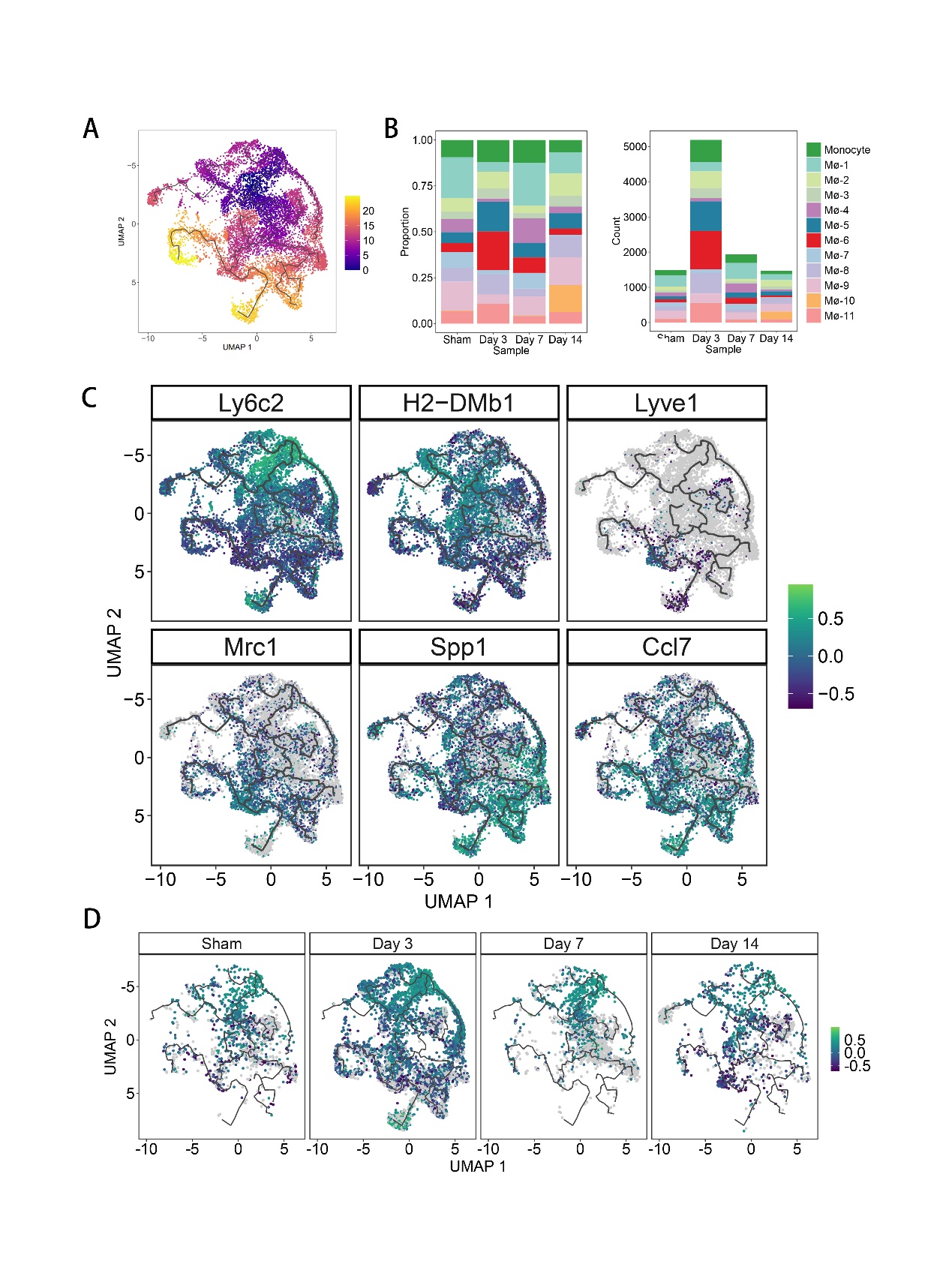

Fig. S6.

(A) Monocle analysis showing the pseudotime of macrophage differentiate trajectory. (B) Fraction of each macrophage subset relative to all cardiac macrophages (Left) and cell number (Right) of each subset. (C) UMAP plot showing the marker gene expression gradients identified within the trajectory. (D) *Ccr2* gene expression gradients identified at 0, 3, 7, and 14 days after MI, showing that some cells within the “*Ccr2*^low^ subset” exhibited high expression level of *Ccr2* on day 3.

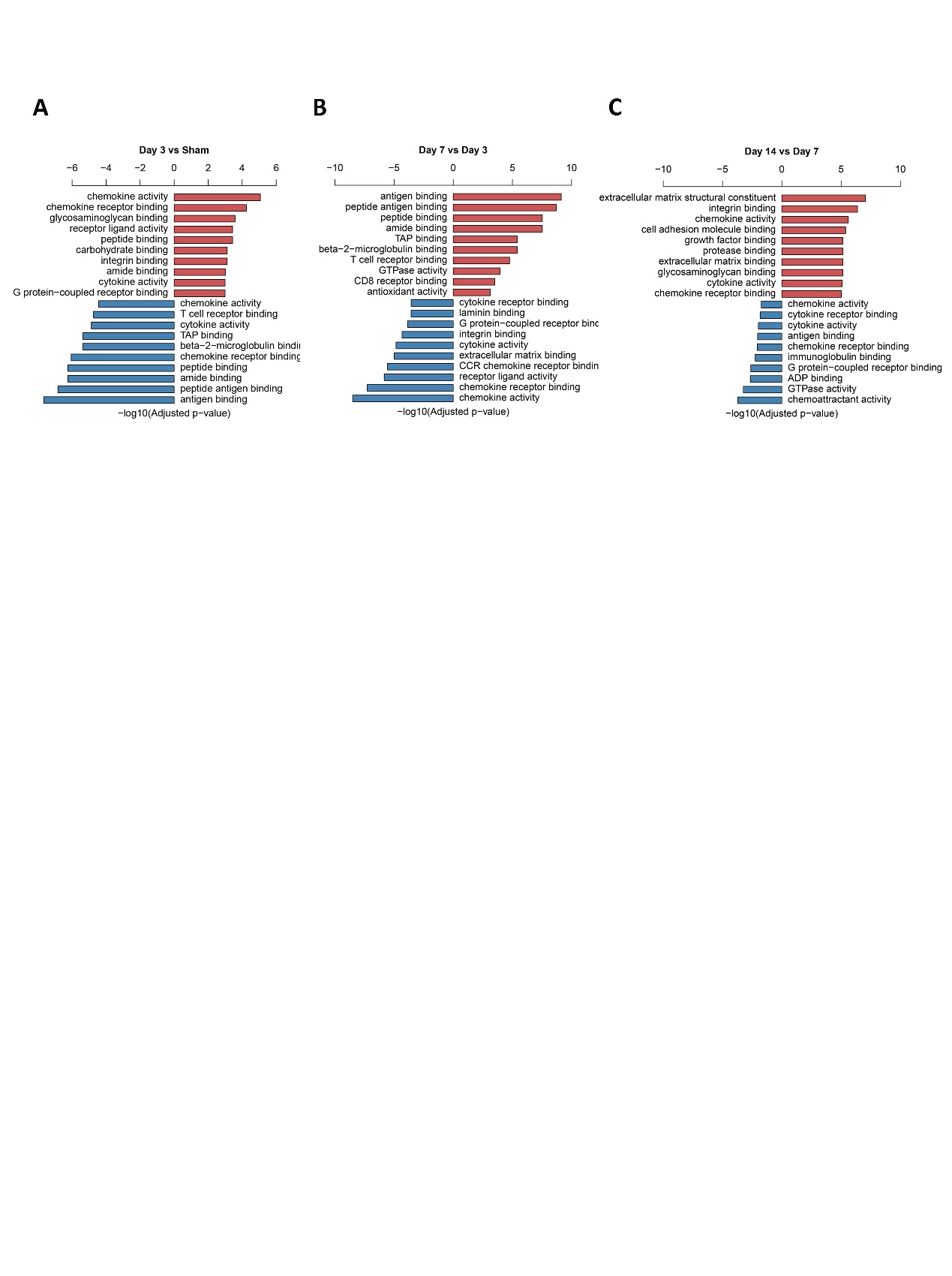

Fig. S7.

GO enrichment terms of differential expression genes in 3d MI and sham (Left), 7d MI and 3d MI (Middle) as well as 14d MI and 7d MI (Right).

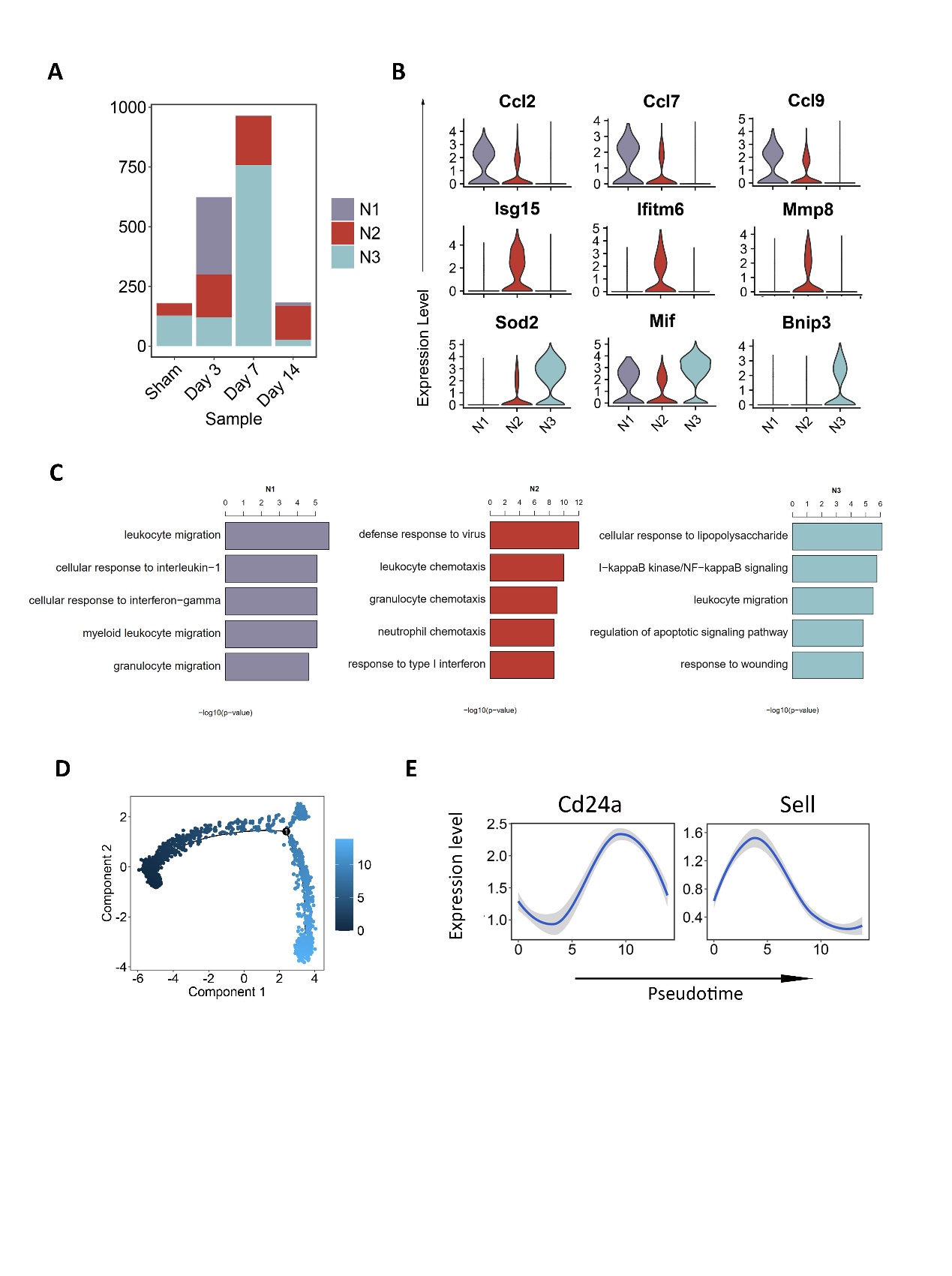

Fig. S8.

(A) Cell number alteration in each neutrophil subset. (B) Expression of feature genes in each neutrophil subset as shown in Fig. 5A. (C) GO enrichment terms of different neutrophil subsets. (D) Monocle analysis showing the pseudotime of neutrophil aging trajectory. (E) Expression of age-related gene were plotted along the pseudotime.

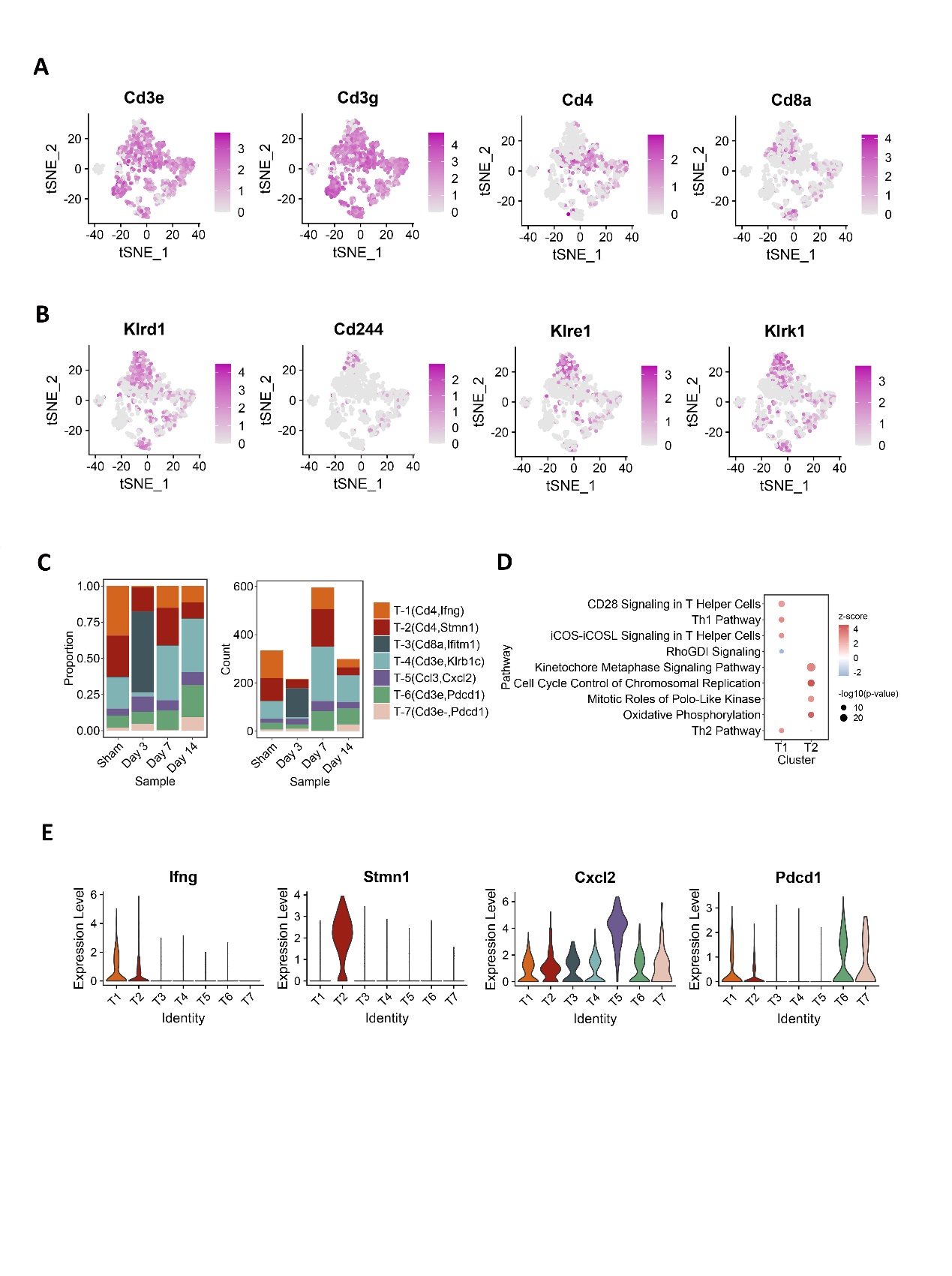

Fig. S9.

Feature plot showing the T cell marker (A) and the NK cell marker (B) expression patterns in cardiac T cell population. (C) Fraction of each T cell subset relative to all cardiac T cells (Left) and cell number (Right) of each subset. (D) Dot plot showing the enriched pathways in Ingenuity-pathway-analysis (IPA) of T1 and T2 Cd4 T cell subsets. *P* values were indicated by circle size and Z scores were indicated by color. (E) Expression of feature genes in each subset as shown in Fig. 5G.

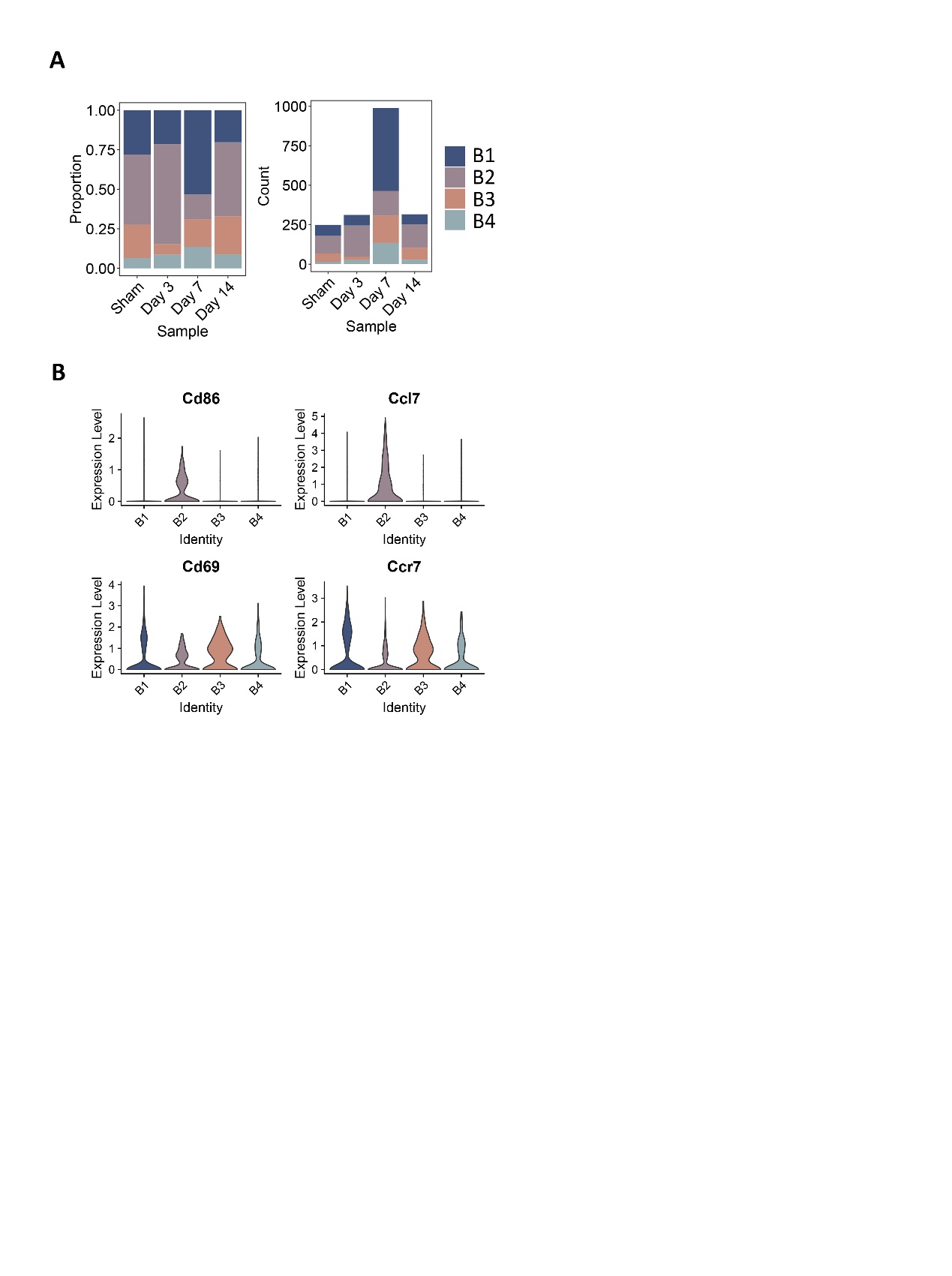

Fig. S10.

(A) Fraction of each B cell subset relative to all cardiac B cells (Left) and cell number (Right) of each subset. (B) Expression of feature genes in each subset as shown in Fig. 5H.

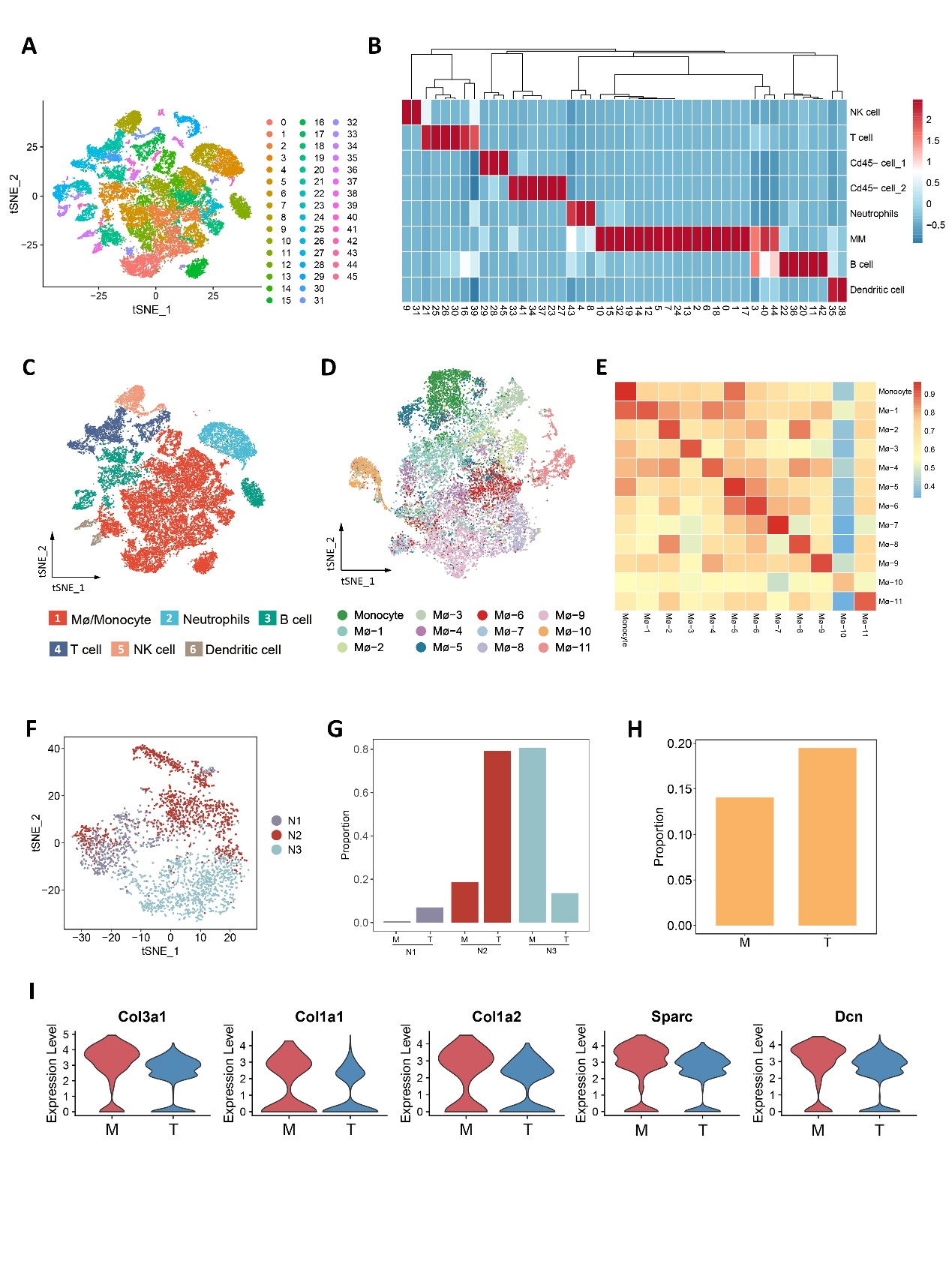

Fig. S11.

(A) tSNE plot showing the unsupervised clustering of the integrated dataset. (B) Heatmap showing the cell type annotation results using Single R. (C) tSNE plot of the integrated dataset from seven samples showing the identical immune cell populations. (D) tSNE plot showing twelve subsets of the Mø/monocyte population from the integrated dataset identified by ItClust. (E) Heatmap showing Spearman correlation of the gene expression detected among macrophage subsets from the MI dataset and the integrated dataset. (F) tSNE plot of three neutrophil subsets from the integrated dataset identified by ItClust. (G) Proportion alteration of neutrophil subsets on day 7 after Tanshinone IIA treatment. (H) Fraction of Mø-10 relative to all macrophages before and after Tanshinone IIA treatment on day 14. (I) Collagen-related gene expression in Mø-10 before and after Tanshinone IIA treatment.

| Estimates | Sham | | | | 3d MI | 7d MI | | 14d MI | |
| --- | --- | --- | --- | --- | --- | --- | --- | --- | --- |
| Estimated Number of Cells | | 2962 | 7506 | | | 5482 | 3942 | | |
| Mean Reads per Cell | | 186509 | | 66215 | | 106104 | | | 159174 |
| Median Genes per Cell | | 1873 | 2413 | | | 1416 | 1645 | | |

Table S1.

Cell Ranger mapping statistics of the MI dataset

| **Cell lineages** | **Marker genes** | | **References** |
| --- | --- | --- | --- |
| Macrophage | Adgre1 Csf1r Cd68 Fcgr1 Lgals3 Arg1 Mrc1 C1qa C1qb | Single-Cell RNA-Seq Reveals the Transcriptional Landscape and Heterogeneity of Aortic Macrophages in Murine Atherosclerosis | |
|  |  | Immune cell census in murine atherosclerosis: cytometry by time of flight illuminates vascular myeloid cell diversity | |
|  |  | Single-Cell Sequencing of the Healthy and Diseased Heart Reveals Cytoskeleton-Associated Protein 4 as a New Modulator of Fibroblasts Activation. | |
|  |  | Apoptosis inhibitor of macrophage depletion decreased M1 macrophage accumulation and the incidence of cardiac rupture after myocardial infarction in mice. | |
|  |  | Atlas of the Immune Cell Repertoire in Mouse Atherosclerosis Defined by Single-Cell RNA-Sequencing and Mass Cytometry | |
| Monocyte | Ly6c2 Ccr2 | Single-Cell RNA-Seq Reveals the Transcriptional Landscape and Heterogeneity of Aortic Macrophages in Murine Atherosclerosis | |
| B cell | Cd79a Cd79b Ly6d Cd19 | Single-Cell RNA-Seq Reveals the Transcriptional Landscape and Heterogeneity of Aortic Macrophages in Murine Atherosclerosis | |
|  |  | Ontogeny of arterial macrophages defines their functions in homeostasis and inflammation | |
| T cell | Cd3e Cd3g Cxcr6 | Atlas of the Immune Cell Repertoire in Mouse Atherosclerosis Defined by Single-Cell RNA-Sequencing and Mass Cytometry | |
|  |  | Single-Cell RNA-Seq Reveals the Transcriptional Landscape and Heterogeneity of Aortic Macrophages in Murine Atherosclerosis. | |
| NK cell | Klra8 Klrb1c Klrc1 Ncr1 Klra4 Klrc2 Klrd1 | Single-Cell RNA-Seq Reveals the Transcriptional Landscape and Heterogeneity of Aortic Macrophages in Murine Atherosclerosis. | |
|  |  | Atlas of the Immune Cell Repertoire in Mouse Atherosclerosis Defined by Single-Cell RNA-Sequencing and Mass Cytometry | |
|  |  | Single-cell transcriptomics of the mouse kidney reveals potential cellular targets of kidney disease. | |
| Dendritic cell | Ccl22 Fscn1 Siglech Tcf4 | Dissecting Immune Circuits by Linking CRISPR-Pooled Screens with Single-Cell RNA-Seq | |
|  |  | A regulatory dendritic cell signature correlates with the clinical efficacy of allergen-specific sublingual immunotherapy | |
|  |  | Atlas of the Immune Cell Repertoire in Mouse Atherosclerosis Defined by Single-Cell RNA-Sequencing and Mass Cytometry | |

Table S2.

Cell markers for all immune cell populations

| Estimates | 3d Tan IIA | | 7d Tan IIA | | 14d Tan IIA |
| --- | --- | --- | --- | --- | --- |
| Estimated Number of Cells | 3928 | 6570 | | | 7674 |
| Mean Reads per Cell | 137037 | | 82255 | 72420 | |
| Median Genes per Cell | 1225 | | 1729 | 1407 | |

Table S3.

Cell Ranger mapping statistics of Tan IIA groups
